# Human Fronto-Striatal Connectivity is Organized into Discrete Functional Subnetworks

**DOI:** 10.1101/2021.04.12.439415

**Authors:** Evan M. Gordon, Timothy O. Laumann, Scott Marek, Dillan J. Newbold, Jacqueline M. Hampton, Nicole A. Seider, David F. Montez, Ashley M. Nielsen, Andrew N. Van, Annie Zheng, Ryland Miller, Joshua S. Siegel, Benjamin P. Kay, Abraham Z. Snyder, Deanna J. Greene, Bradley L. Schlaggar, Steven E. Petersen, Steven M. Nelson, Nico U.F. Dosenbach

**Author notes:** Correspondence (E.M.G.).

## Abstract

The striatum is interconnected with the cerebral cortex via multiple recurrent loops that play a major role in many neuropsychiatric conditions. Primate cortico-striatal connections can be precisely mapped using invasive tract-tracing. However, noninvasive human research has not mapped these connections with anatomical precision, limited by the practice of averaging neuroimaging data across individuals. Here we utilized highly-sampled resting-state functional connectivity MRI for individually-specific precision functional mapping of cortico-striatal connections. We identified ten discrete, individual-specific subnetworks linking cortex—predominately frontal cortex—to striatum. These subnetworks included previously unknown striatal connections to the human language network. The discrete subnetworks formed a stepped rostral-caudal gradient progressing from nucleus accumbens to posterior putamen; this organization was strongest for projections from medial frontal cortex. The stepped gradient organization fit patterns of fronto-striatal connections better than a smooth, continuous gradient. Thus, precision subnetworks identify detailed, individual-specific stepped gradients of cortico-striatal connectivity that include human-specific language networks.

## INTRODUCTION

The striatum (putamen, caudate, nucleus accumbens, and globus pallidus) is critically important for optimizing and executing goal-directed behaviors across multiple modalities (Cohen and Frank, 2009; Haber, 2016), ranging from motor planning and control (Grillner et al., 2005), to manipulation of items in working memory (Frank et al., 2001), to complex value judgments (Schultz et al., 1998; Schultz and Dickinson, 2000). The striatum works in close concert with the cortex—and especially the frontal cortex—via a series of parallel, partially segregated cortico-striato-thalamo-cortical circuits (Haber, 2003; Nakano et al., 2000). The integrity of cortico-striatal connectivity is critical for healthy brain function, as its disruption causes devastating neurological and psychiatric symptoms in disorders such as Parkinson’s and Huntington’s disease (Blumenstock and Dudanova, 2020; Bunner and Rebec, 2016). Even relatively minor alterations in cortico-striatal circuitry are thought to play a major role in symptoms observed in Tourette syndrome (Greene et al., 2017; Mink, 2001), schizophrenia (Simpson et al., 2010), obsessive-compulsive disorder (Harrison et al., 2009), depression (Borsini et al., 2020), and autism (Li and Pozzo-Miller, 2019). However, cortico-striatal circuits are some of the least well-understood pathways in the human brain, as their striatal elements are small and exceedingly difficult to dissociate from each other using non-invasive imaging. Precise, detailed, individual-specific characterization of cortico-striatal circuits is a fundamental first step to understanding how they help generate complex behaviors, and is vitally important for treating many neurodegenerative and neuropsychiatric disorders, especially with targeted neurostimulation (e.g., using deep brain stimulation [DBS], or focused ultrasound).

Non-human primate research using autoradiographic tracer injections has demonstrated that the striatum receives topographically ordered projections from a wide variety of cortical sources (Cavada and Goldman-Rakic, 1991; Cheng et al., 1997; Chikama et al., 1997; Selemon and Goldman-Rakic, 1988, 1985; Steele and Weller, 1993; Van Hoesen et al., 1981; Webster et al., 1993; Yeterian and Pandya, 1998, 1993; Yeterian and Van Hoesen, 1978), though projections from frontal cortex are thought to be the main input driving striatal function (Haber, 2003). Non-human primate fronto-striatal circuits have an ordered organization that is mirrored in striatum and cerebral cortex (Haber, 2016). Specifically, rostral portions of frontal cortex project to punctate regions within rostral striatum (Haber et al., 1995; Kunishio and Haber, 1994), while progressively more caudal frontal cortex projects to progressively caudal striatal targets (Averbeck et al., 2014; Calzavara et al., 2007; Flaherty and Graybiel, 1994, p. 1975; Künzle, 1975; Selemon and Goldman-Rakic, 1985).

The fronto-striatal organization observed in non-human primates is largely assumed to have been conserved in the human brain (Haber, 2016). However, the complex cognitive and psychiatric symptoms caused by striatal dysfunction point towards human-specific aspects of fronto-striatal connectivity. Further, relative to non-human primates, the prefrontal cortex is expanded in humans (Smaers et al., 2011), and pathways for language processing have developed (Dick and Tremblay, 2012; Friederici, 2017). It is unknown how human-specific circuits integrate into fronto-striatal organization.

Precisely mapping human fronto-striatal connections with non-invasive imaging has been challenging. Techniques such as diffuse optical tomography, electroencephalography, and magnetoencephalography cannot obtain signal from the brain center. MRI-based techniques such as resting-state functional connectivity (RSFC), which involves identifying correlated patterns of activity in spatially distant regions of the brain (Biswal et al., 1995), have proven successful in describing the brain’s network organization (Power et al., 2011; Yeo et al., 2011). However, these techniques suffer rapid drop-off of signal-to-noise (SNR) ratios as distance increases from the head coil. Standard approaches for overcoming low SNR in the brain center require registering study participants to a common atlas and averaging their data. Such group-averaged neuroimaging studies have been limited to describing broad, nonspecific connections between large swaths of striatum and widespread regions of cortex, distributed large-scale networks, or entire cortical lobes (Badre and Frank, 2012; Choi et al., 2012; Di Martino et al., 2008; Draganski et al., 2008; Greene et al., 2014; Jarbo and Verstynen, 2015; Jeon et al., 2014; Marquand et al., 2017; Mestres-Missé et al., 2012; Morris et al., 2016; Parkes et al., 2017; Tziortzi et al., 2014; Verstynen et al., 2012).

There is significant cross-individual spatial variability in the network organization of both the cortex (Braga and Buckner, 2017; Gordon et al., 2017b, 2017c; Harrison et al., 2015; Li et al., 2019; Wang et al., 2015) and the thalamus (Greene et al., 2020), such that features of functional brain organization are not present at the same location in all subjects. Consequently, group averaging neuroimaging approaches fail to identify spatially localized organizational features that can be observed in individuals (Bijsterbosch et al., 2018; Feilong et al., 2018; Gordon et al., 2017a; Harrison et al., 2020), and this issue is likely magnified in the striatum due to its small size. Thus, prior use of group-averaged data may have obscured important aspects of fronto-striatal organization present in individual humans.

Recent approaches for noninvasively describing the individual-specific organization of the brain using RSFC (termed precision functional mapping, or PFM), have delineated the large-scale network-level organization of the cerebral cortex (Braga et al., 2020; Braga and Buckner, 2017; Gordon et al., 2017c; Laumann et al., 2015) and thalamus (Greene et al., 2020) in individuals. We recently extended PFM to demonstrate the presence of subnetwork structures within classic large-scale networks that comprise strong, functionally differentiated connections between spatially specific cortical and subcortical regions (Gordon et al., 2020). The high spatial specificity of connections that can be identified using this subnetwork-PFM technique makes it ideal for characterizing precise, spatially-specific fronto-striatal connections.

Precise characterization of individual-specific human cortico-striatal organization would allow us to address fundamental questions in neuroscience, such as “is brain organization continuous, or discrete?” The rostral-caudal organization of fronto-striatal projections has been termed a “functional gradient” (Haber, 2003), in that it describes a progression along a single rostral-caudal axis. Group-averaged neuroimaging data have been used to present functional cortico-striatal gradients as a continuous transition along a primary organizational axis (Jarbo and Verstynen, 2015; Marquand et al., 2017; Mestres-Missé et al., 2012; O’Rawe et al., 2019; Raut et al., 2020; Vogelsang and D’Esposito, 2018). However, the conceptualization of a continuous gradient organization of the brain must also be reconciled with the long-held perspective that the cortex is organized into discrete, noncontinuous areas that are networked together (Amunts and Zilles, 2015; Brodmann, 1909; O’Leary et al., 2007; Sejnowski and Churchland, 1989). Within these areas, functional, connectional, topographic, and architectonic properties are relatively constant, while borders between areas exhibit abrupt, discontinuous changes in these properties (Cohen et al., 2008; Felleman and Van Essen, 1991). The question of whether brain organization is better conceptualized as continuous gradients or discontinuous areas is a critical one, as conceptual models can strongly influence experimental and analytical approaches. For example, when studying a neuropsychiatric disorder such as schizophrenia, should the focus be on location, size, function, and connections of discrete cortical and striatal areas? Or are neuropsychiatric disorders better captured by alterations in the endpoints, direction, shape, and descent steepness of continuous cortical and striatal gradients?

Here, we precisely characterized fronto-striatal connections in individual humans. We identified subnetworks that describe cortico-striatal connections in ten individual human brains using >5 hours of RSFC data per-individual (Midnight Scan Club [MSC] dataset). We examined the degree to which the observed organization of cortico-striatal connections converged or differed from existing models of fronto-striatal connectivity based on non-human primate data (Haber, 2016). We then tested whether individual-specific fronto-striatal connections were best represented as a smooth, continuous progression, or whether they were best represented as discontinuous functional areas.

## RESULTS

### Striatal Subnetworks are Individually Specific but Spatially Similar Across Subjects

We delineated individual-specific subnetworks connecting striatum to cortex and matched them across subjects following procedures described in (Gordon et al., 2020). Briefly, in each subject we calculated RSFC strength between each pair of points in the brain, and then used a data-driven subnetwork detection procedure designed to identify and operate upon the strongest functional connections of every point in the brain. We then matched subnetworks with striatal and cortical representation across subjects. See Methods for details.

Ten separate subnetworks with striatal and cortical representation were identified that were spatially and topologically consistent across subjects, with some inter-individual variability in the localization of each subnetwork. See Fig 1 for an example subject; see Fig S1 for all subjects; see Fig 2 for the overlap of each subnetwork across subjects; and see Table S1 for information about the consistency of connections. These ten subnetworks existed as substructures within several known large-scale brain networks, including the Default-Mode, Salience, Cingulo-opercular, Fronto-parietal, Language, and Somatomotor networks (see Fig 3 for an example subject; see Table S1 for mode network memberships across subjects). The spatial representations of these subnetworks spanned from posterior putamen to nucleus accumbens in striatum, and from central sulcus and parietal/temporal lobes in cortex to subgenual cingulate/orbitofrontal cortex.

**Figure 1:**
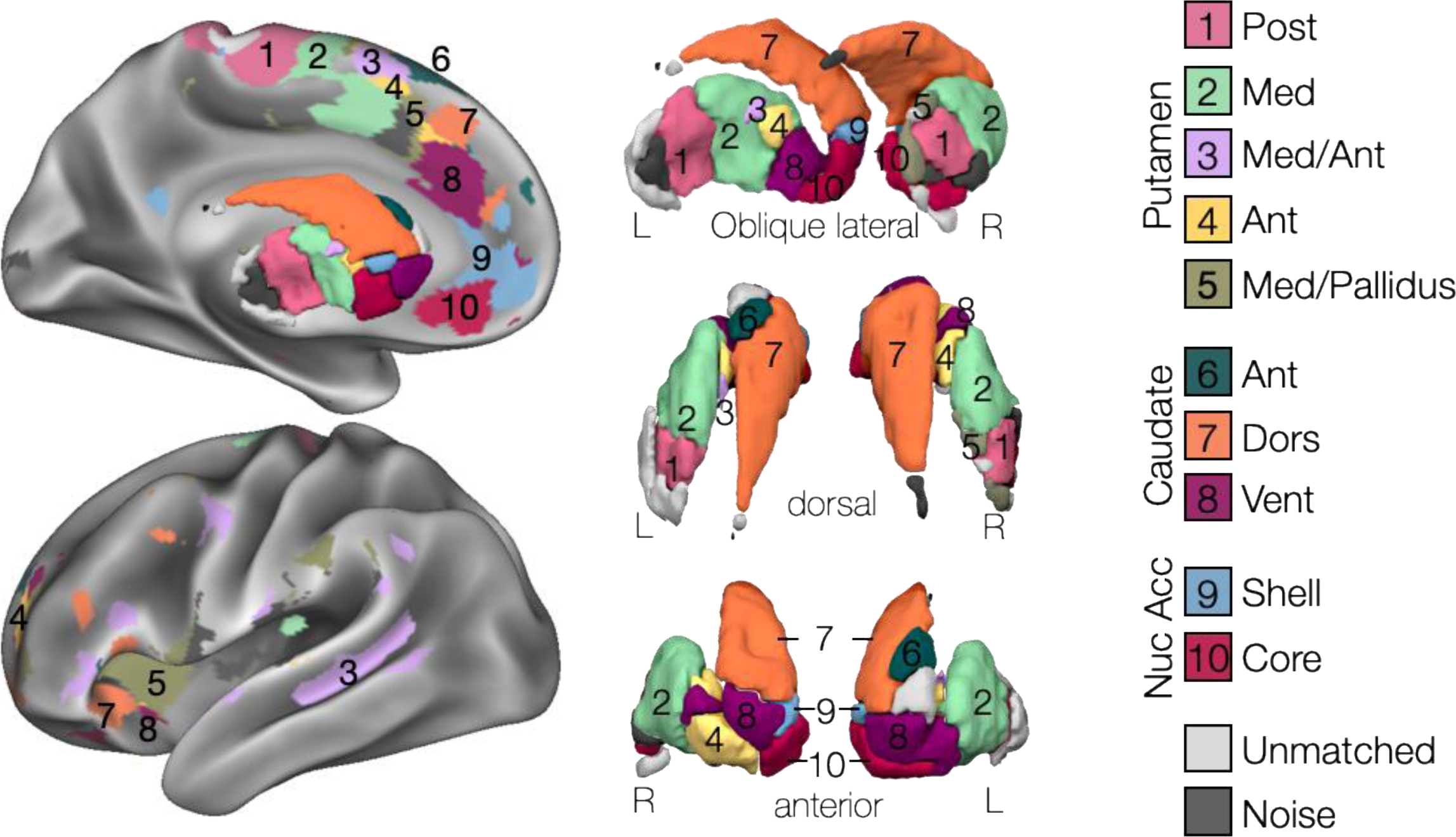
Cortico-Striatal Subnetworks in an Exemplar Subject (MSC01). Left: Subnetwork representation in left hemisphere cortex and striatum. Right: Subnetwork representation in bilateral striatum from dorsal (top), anterior (middle), and oblique lateral-posterior (bottom) views.

**Figure 2:**
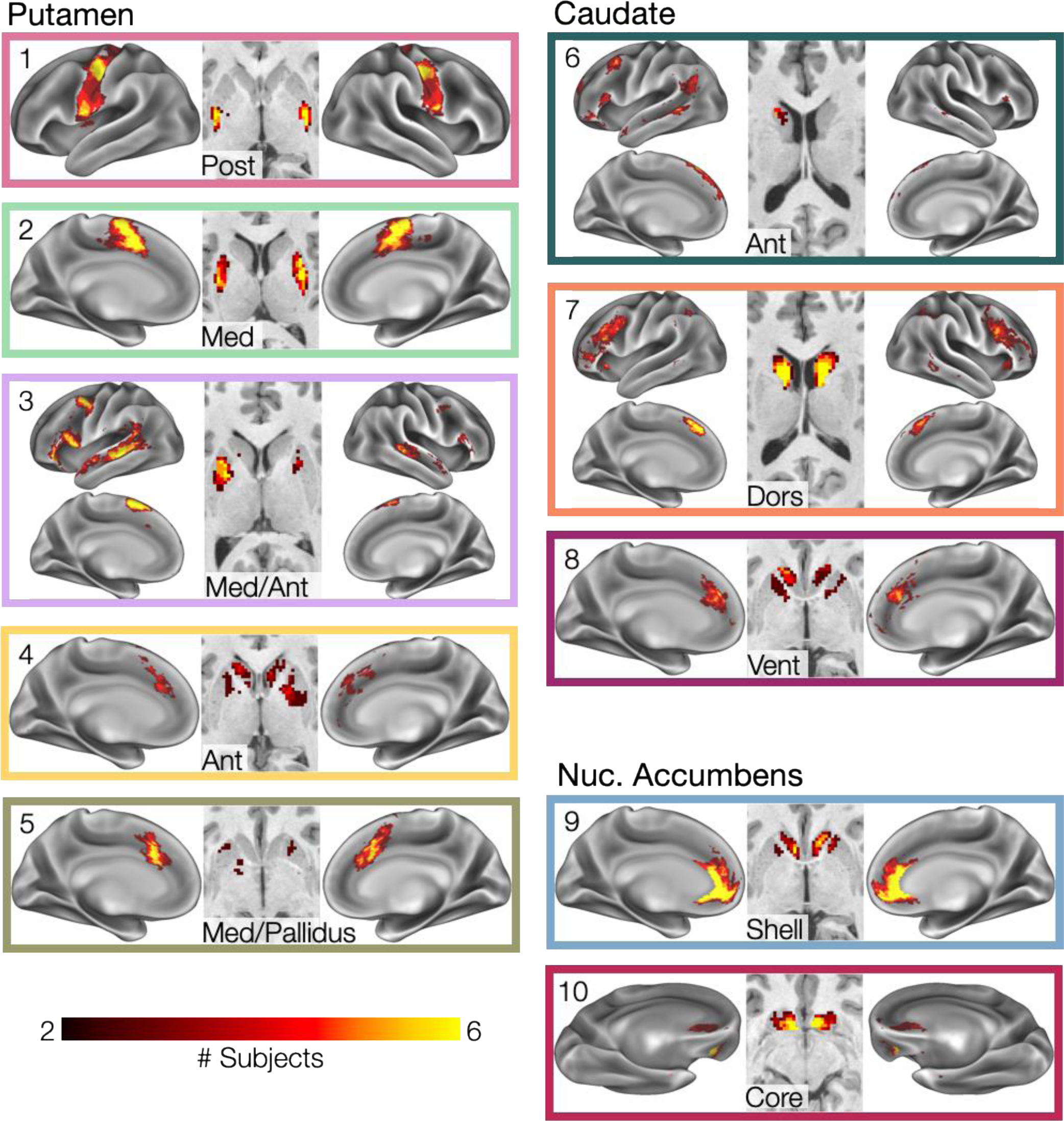
Striatal Subnetwork Overlap across Subjects. Cross-subject overlap of matched striatal subnetworks. Heat map indicates number of subjects with spatial overlap of matched subnetworks at each point in the brain.

**Figure 3:**
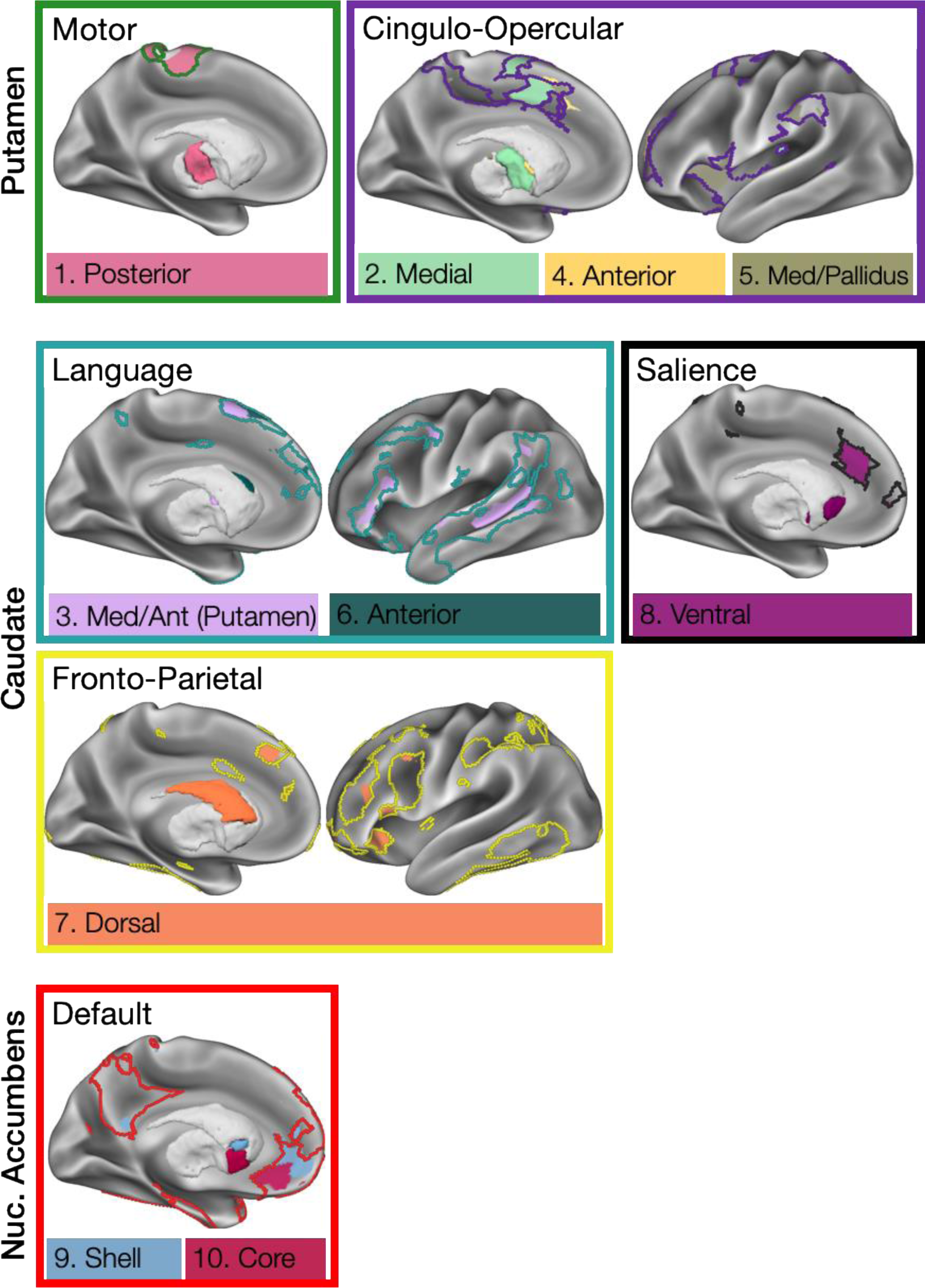
Striatal Subnetworks Are Substructures within Canonical Functional Networks. Striatal subnetworks in an exemplar subject (MSC01) are contained within known large-scale cortical networks (colored borders).

Across subjects, an average of 95% (range: 91% to 99%) of striatal voxels were matched to one of the ten subnetworks. Thus, the subnetworks described here represent the large majority of strong functional cortico-striatal connections, with very little of the striatum containing idiosyncratic subnetwork connections that cannot be compared across subjects.

These subnetworks showed significant overlap with the non-human primate-based model of cortico-striatal connections advanced by (Haber, 2016) (Fig 4). First, these subnetworks, which represent very strong cortico-striatal functional connections, most commonly connected striatum to frontal cortex. In each subject, we calculated the percent of cortical vertices with any subnetwork representation that were within each cortical lobe (Fig S2A). Across subjects, we found that 70% ± 6% (mean ± SD) of these subnetworks’ cortical vertices were within frontal lobe, which was higher in every subject than in the insula (10% ± 2%), parietal lobe (9% ± 3%), temporal lobe (10% ± 6%), or occipital lobe (1% ± 1%); all paired ts(9) > 17.0, all ps < 10^-7^, indicating that subnetwork connections were most commonly to frontal lobe. Similarly, we evaluated the strength of within-subnetwork cortico-striatal RSFC for each lobe in each subject (Fig S2B). Across subjects, within-subnetwork RSFC between striatal and frontal regions was 0.25 ± .02 (mean ± SD), which was higher in every subject than RSFC between striatum and insula (0.15 ± 0.03), parietal lobe (0.11 ± 0.04), temporal lobe (0.10 ± 0.02), or occipital lobe (0.12 ± 0.06); all paired ts(9) > 6.8, all ps < 0.0002, indicating that the few non-frontal subnetwork connections that were identified were weaker than frontal connections.

**Figure 4:**
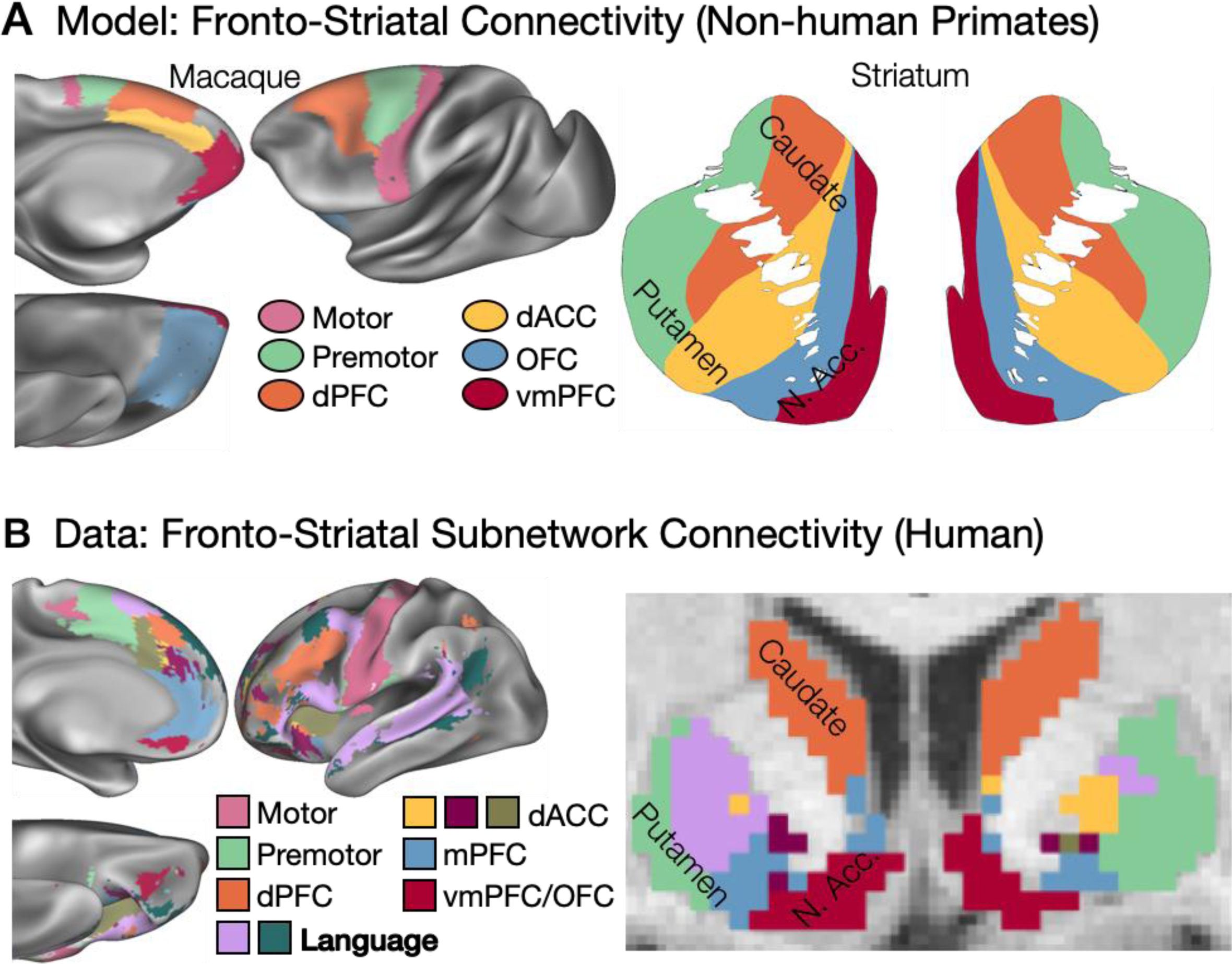
Non-Human Primate Model and Actual Organization of Human Cortico-Striatal Connectivity. A) Prior organizational model of fronto-striatal connections in cortex (left) and striatum (right), based primarily on tract-tracing work in non-human primates, proposed by Haber, 2016. Images are adapted from Haber, 2016, with anatomical locations displayed on a macaque cortex. B) Organization of cortico-striatal connections observed in the present work. Cortical vertices/voxels are colored based on the most common subnetwork present across the ten subjects. A putative Language network (bolded in the inset legend) was not identified in the Haber model. Note that a Motor subnetwork is present posterior of the displayed slice in putamen, and a second Language network is present anterior of the displayed slice in left caudate.

Second, the spatial distributions of these subnetworks in large parts converged with the Haber (2016) model (Fig 4), with several important exceptions (see below). Projections to posterior and middle putamen originated in motor and premotor cortex, respectively; projections to dorsal caudate originated from dorsomedial and dorsolateral prefrontal cortex; and projections to ventral anterior caudate and medial putamen originated from the dorsal anterior cingulate cortex. Projections to nucleus accumbens originated from medial and orbitofrontal cortex; though notably, the targets of the medial and orbitofrontal cortex projections within nucleus accumbens (to the shell and core, respectively) appeared reversed relative to the non-human primate model.

Importantly, we discovered two subnetworks missing from the non-human primate-derived model of cortico-striatal connections advanced by (Haber, 2016) (Fig 4): 1) a subnetwork (Figs 1-3, #3, Med Ant Putamen, pink) with representation primarily in medial putamen and sometimes lateral caudate head, as well as in a distributed set of cortical regions including dorsomedial prefrontal cortex, inferior frontal gyrus, posterior middle frontal gyrus, and superior temporal sulcus; and 2) a subnetwork (Figs 1-3, #6, Ant Caudate, dark green) with representation in anterior lateral caudate and distributed cortical regions immediately adjacent to the medial putamen subnetwork. See Fig S3A for these subnetworks in all subjects. Notably, the cortical distributions of these subnetworks— especially the medial putamen subnetwork—converge with the known distribution of the human language network (Braga et al., 2020).

Like the human language network, these two subnetworks were left-lateralized, with more extensive left than right hemisphere representation in both cortical and subcortical structures. To quantify this observation, for each subnetwork we calculated a laterality index as ([# left hemisphere vertices/voxels] – [# right hemisphere vertices/voxels]) / [# total vertices/voxels], separately in cortex and in striatum. With this index, positive values indicate left lateralization, while negative values indicate right lateralization. We found that, in every subject, these two subnetworks were left-lateralized in both cortex and striatum (medial putamen subnetwork: cortex mean laterality=.62±.24, one-sample t(9)=7.9, p<10^-4^; striatum mean laterality=.73±.32, t(9)=7.2, p<10^-4^; lateral caudate subnetwork: cortex mean laterality=.67±.34, one-sample t(9)=5.7, p<0.001; striatum mean laterality=.72±.36, t(9)=5.4, p<0.002); see Fig S3B. By contrast, no other subnetwork exhibited consistent lateralization across subjects (all ts(9)<2.2, all ps>0.05 in both cortex and striatum).

### Fronto-Striatal Organization is Better Explained as a Stepped than a Continuous Gradient

Fronto-striatal connectivity is often presented as exhibiting a smooth and continuous rostral-caudal gradient (e.g., model in Fig 5A, left). Visual examination of the subnetworks identified here (see Fig 1) does support the presence of a rostral-caudal gradient; however, it also suggests that this gradient may be not smooth, but comprised of a progression of discontinuous cortical areas with discrete connectivity profiles (model in Fig 5A, right). Such a gradient may be conceptualized as “stepped” rather than smooth and continuous. A stepped gradient would be consistent with both the postulated rostral-caudal gradient of fronto-striatal projections (Vogelsang and D’Esposito, 2018) as well as with the areal model of cortex (Brodmann, 1909; Churchland and Sejnowski, 1988). As a third possibility, both the continuous and stepped models could simultaneously be present, such that regions exhibit categorically distinct fronto-striatal connections from each other, but a rostral-caudal gradient also exists overlaid on top of the discrete regions. Here, we used linear regression models to test whether the locations of strong fronto-striatal connections were best explained by continuous rostral-caudal position, or by discrete subnetwork identities, or by both.

**Figure 5:**
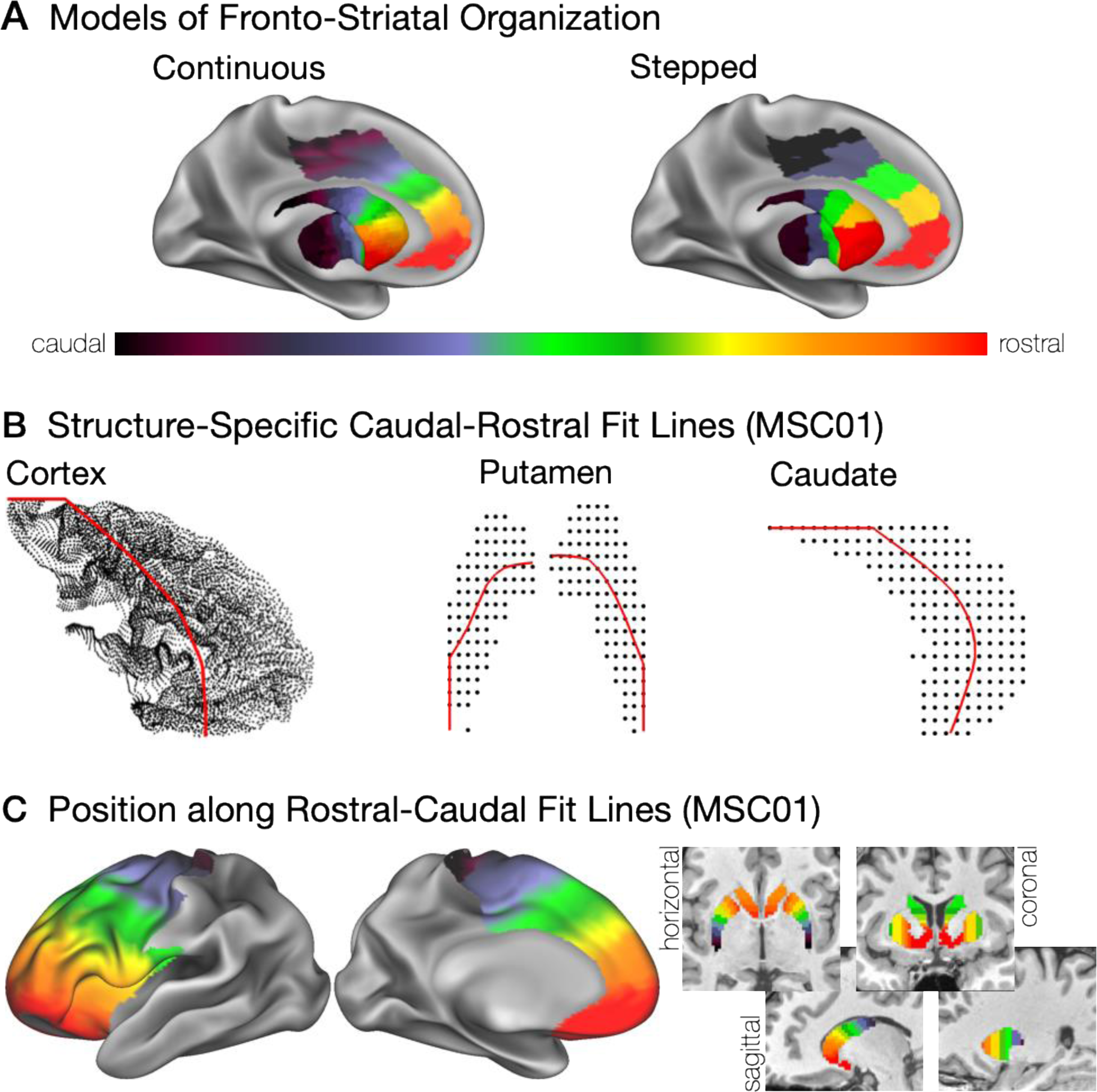
Modeling Fronto-Striatal Gradient Organization: Continuous vs Stepped. A) Conceptually, fronto-striatal connections may be organized as a continuous (left) or a stepped (right) mirrored rostral-caudal gradient. B) Rostral-caudal orientation fit lines are shown for example subject MSC01 within frontal cortex plus anterior insula (left), left and right putamen (middle), and caudate (right). C) Position along the rostral-caudal fit lines is shown for example subject MSC01 for all points within frontal cortex plus anterior insula (left) and striatum (right). These rostral-caudal positions were used to compare models of organization described in (A). Note that while striatal connections with non-frontal sources were observed (Figs 1-3, Fig S2), rostral-caudal modeling here is restricted to the frontal cortex.

We first had to define the rostral-caudal position for each brain location of interest. We defined a separate rostral-caudal axis for the caudate, the left and right putamen, and the prefrontal cortex (Fig 5B), each with its own unique geometry. We then labeled each striatal voxel and each frontal cortex vertex according to its position along that fit line (i.e., the point along the structure-specific fit line that is closest to that voxel / vertex). The calculated rostral-caudal position of each voxel / vertex can be seen in Fig 5C. Note that we elected to use this anatomically-based rostral-caudal orientation rather than explicitly fitting a gradient to the RSFC data because we did not want the continuous gradient overfit to the data when comparing against a discrete organizational model.

We then conducted two separate, symmetric series of regressions in every subject. First, for every frontal/insular cortical vertex with a subnetwork identity, we calculated the voxel within the striatum that had the strongest functional connectivity with that vertex. As stronger functional connectivity is arguably likely to represent more direct connections, we take this voxel of maximal connectivity to be the striatal target of that cortical vertex.

To test the continuous gradient hypothesis, which holds that the rostral-caudal positions of connected cortical and subcortical regions should vary continuously with each other, we regressed the rostral-caudal position of the cortical source vertex against the continuous rostral-caudal position of the striatal target voxel. To test the discrete, stepped gradient hypotheses, we conducted a one-way ANOVA examining whether the rostral-caudal position of the striatal target voxel was explained by the subnetwork identity of the cortical source vertex. In each case, the adjusted R^2^ value was calculated as the measure of variance explained, in order to allow direct comparisons between these two regressions which had differing numbers of independent variables. It is important to note that the dependent variable in both of these regressions was the rostral-caudal position of the subcortical target voxel. As a result, the adjusted R^2^ values are directly comparable between the two; and further, this rostral-caudal position is not a priori known from either the cortical vertex’s position or its subnetwork identity. Finally, to test whether both the continuous and discrete models might be simultaneously true, we entered both the cortical position and subnetwork identity into an ANCOVA model testing against subcortical voxel position.

We found that both the continuous and the discrete, stepped gradient hypotheses successfully fit the data, but that the discrete stepped hypothesis explained more variance in the data (Fig 6, A,C,E). Across subjects, a cortical vertex’s rostral-caudal position explained 17% ± 16% (mean ± SD) of the variance in the rostral-caudal position of its connected striatal voxel (see Fig 6A for an example subject; Fig 6E, top for all subjects). By contrast, the vertex’s subnetwork identity explained 56% ± 13% of the variance in the rostral-caudal position of its connected striatal voxel (see Fig 6C for an example subject; Fig 6E, top for all subjects). Subnetwork identity explained more variance than rostral-caudal position in every subject.

**Figure 6:**
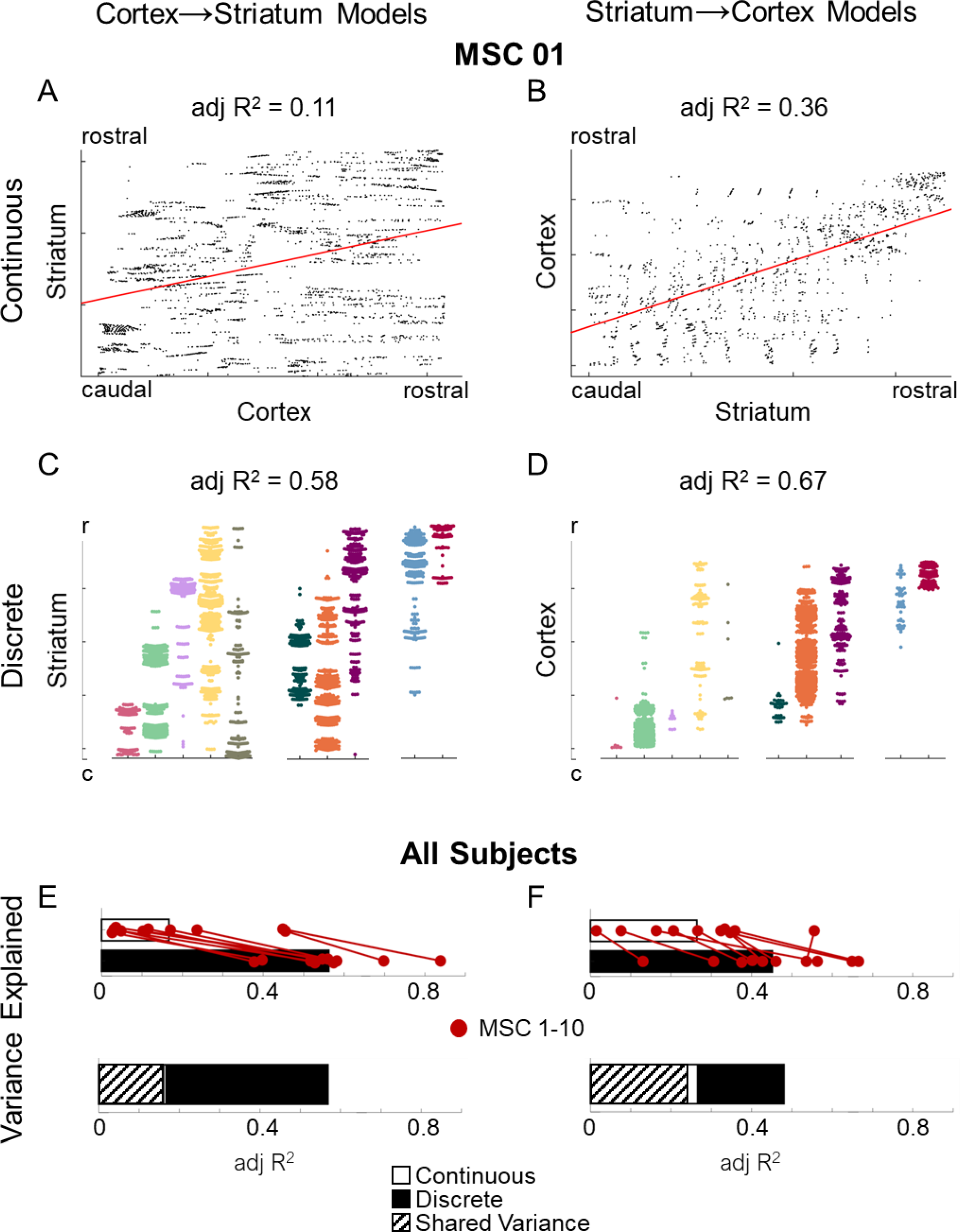
Stepped Gradients Composed of Subnetworks Explain more Variance in Fronto-Striatal Organization than Continuous Gradients. In example subject MSC01, A) the continuous rostral-caudal position of a cortical vertex (x-axis) explained variance in the rostral-caudal position of its most strongly connected striatal target (y-axis), and vice versa (B). However, the discrete subnetwork identity of a cortical vertex (delineated by color and by x-axis position) explained more variance in the rostral-caudal position of its striatal target (y-axis), and vice versa (D). In A-D, each dot represents one cortical vertex (A,C) or striatal voxel (B,D). Across subjects, for both the cortex→striatum regressions (E) and the striatum→cortex regressions (F), both continuous rostral-caudal position and discrete subnetwork identity explained substantial variance, but subnetwork identity explained more variance in every subject (top). When both factors were employed in the same model (bottom), the discrete subnetwork identity still explained substantial unique variance, but the continuous gradient factor explained almost no unique variance.

When both factors were entered into the same ANCOVA model, only the discrete stepped gradient hypothesis explained unique variance in the data (Fig 6E, bottom). Across subjects, the rostral-caudal position of a cortical vertex explained only 0.6% ± 0.8% of unique variance in the position of its striatal target, after controlling for subnetwork identity. By contrast, the subnetwork identity of the cortical vertex still explained 40% ± 7.5% of unique variance in the position of its striatal target, after controlling for rostral-caudal position.

The above regressions were asymmetric: they explained the rostral-caudal position of a subcortical target based on the position and/or subnetwork identity of its cortical source. To ensure symmetry of these effects, we repeated all of the regressions in the reverse direction, in which the most strongly connected cortical source vertex was calculated for each striatal voxel, and the rostral-caudal position of a cortical source vertex was explained based on the position and/or subnetwork identity of the striatal voxel.

In these regressions, once again both the continuous and the discrete, stepped gradient hypotheses successfully fit the data, but the discrete stepped hypothesis explained more variance in the data (Fig 6, B,D,F): a subcortical voxel’s rostral-caudal position explained 25% ± 17% (mean ± SD) of variance in the rostral-caudal position of its connected cortical vertex (see Fig 6B for an example subject; Fig 6F, top for all subjects), but the voxel’s subnetwork identity explained 45% ± 16% of variance (Fig 6D for an example subject; Fig 6F, top for all subjects), with subnetwork identity explaining more variance than rostral-caudal position in 9/10 subjects.

When both factors were entered into one model, again only the discrete stepped gradient hypothesis explained unique variance in the data (Fig 6F, bottom), with the rostral-caudal position of a subcortical voxel explaining only 3% ± 4% of unique variance after controlling for subnetwork identity, and the subnetwork identity of the voxel still explaining 22% ± 12% of unique variance. These findings indicate that the rostral-caudal progression of strong fronto-striatal connections is organized as a stepped gradient composed of a series of discrete subnetworks.

The progression of subnetworks along the rostral-caudal axis is more striking on the medial surface of the frontal cortex than on the lateral surface (e.g. Fig 1). It is possible that the relative explanatory power of continuous vs stepped gradients might vary based on whether projections are from the medial or lateral surface. To explore this possibility, we repeated the regressions above twice: once while restricting cortical source vertices to only the medial, and then to the lateral, frontal cortical surfaces. We found that on the medial surface, adjusted R^2^ values were much higher for both the continuous (cortex→striatum: 33% ± 17%; striatum→cortex: 41% ± 12%) and the stepped (cortex→striatum: 61% ± 13%; striatum→cortex: 66% ± 12%) gradient models, but the stepped model explained more variance than the continuous model in every subject, in both directions. On the lateral surface, adjusted R^2^ values for the continuous gradient (cortex→striatum: 18% ± 24%; striatum→cortex: 24% ± 14%) were similar to those observed across the whole frontal cortex, but adjusted R^2^ values for the stepped gradient were slightly lower (cortex→striatum: 50% ± 13%; striatum→cortex: 38% ± 14%); however, the stepped model explained more variance than the continuous model in every subject in the striatum→cortex direction, and in 9/10 subjects in the cortex→striatum direction. These results suggest that the rostral-caudal organization of fronto-striatal connections, which is organized as a stepped gradient, is more prominent in the medial than the lateral frontal cortex.

## DISCUSSION

In this work, we identified ten subnetworks representing very strong, anatomically precise, individually-specific connections between cortex and striatum that could be matched across subjects. The identification of these connections within individual humans represents a substantial breakthrough in our ability to understand these circuits that are both critical for complex behaviors and implicated in many neuropsychiatric disorders. The neurobiological validity of the observed cortico-striatal subnetwork structures is supported by their general convergence with findings from invasive, anatomically precise tract-tracing studies in non-human primates. Critically, the identified subnetworks also diverge from non-human primate-based models of fronto-striatal connectivity in several ways, most prominently by identifying left-lateralized striatal elements of the human language network.

### Most Cortico-striatal Subnetworks Converge with Primate Tract-tracing

We identified striatal subnetworks composed of only very strong functional connections in single individuals, and we found that 1) the majority of these very strong cortico-striatal connections were from frontal cortex, and 2) frontal regions of these subnetworks had stronger connectivity with striatum than non-frontal regions, suggesting that the non-frontal regions we did observe were less strongly associated with striatal activity. This converges with non-human primate work arguing that, while striatum does receive inputs from diverse cortical sources across frontal, parietal, insular, and temporal cortex (Cavada and Goldman-Rakic, 1991; Cheng et al., 1997; Chikama et al., 1997; Selemon and Goldman-Rakic, 1988, 1985; Steele and Weller, 1993; Van Hoesen et al., 1981; Webster et al., 1993; Yeterian and Pandya, 1998, 1993; Yeterian and Van Hoesen, 1978), the strongest, most driving projections to striatum are argued to arise from frontal lobe (Haber, 2003). Thus, the present subnetwork connections represent a substantial increase in specificity over prior human neuroimaging work that identified cortico-striatal connections originating from large swaths of cortex and from distributed networks across multiple cortical lobes (Choi et al., 2012; Greene et al., 2014; Morris et al., 2016; Parkes et al., 2017; Tziortzi et al., 2014).

The identified fronto-striatal circuits were highly ordered as a topographically mirrored rostral-caudal gradient of cortico-striatal projections. This organization converged with the organization of these circuits proposed based on findings from non-human primates (Haber, 2016, 2003); see Fig 4. Specifically, moving from caudal to rostral striatum:

1. We found functional connections between somatomotor regions in the central sulcus and the posterior putamen (Fig 4, magenta), consistent with previous observations of projections from precentral motor cortex to posterior putamen in humans (Zeharia et al., 2015) and non-human primates (Flaherty and Graybiel, 1994; Künzle, 1975). These connections are likely related to striatal control over motor function.
2. We found functional connections between regions just anterior to the somatomotor strip, particularly in dorsomedial prefrontal cortex, and the middle lateral putamen (Fig 4, green), consistent with previous observations in non-human primates of projections from pre-/supplementary motor cortex to middle lateral putamen (Haber, 2003; Künzle, 1978). These circuits are engaged during performance of complex (non-automatized) motor skills (Boecker et al., 1998).
3. We found functional connections between dorsomedial and dorsolateral prefrontal cortex, within the fronto-parietal network, and a large extent of dorsal caudate (Fig 4, orange). This observation is consistent with previous accounts of projections from dorsolateral prefrontal cortex to terminals in dorsal caudate that span a significant rostral-caudal extent (Arikuni and Kubota, 1986; Haber et al., 2006; Selemon and Goldman-Rakic, 1985), which have been argued to play a critical role in working memory (Partiot et al., 1996).
4. We found several sets of functional connections between dorsal anterior cingulate cortex / anterior insula and the anterior caudate/putamen (Fig 4, yellow / maroon / brown). These observations align with those from previous investigations of projections from primate dorsal anterior cingulate to anterior caudate and putamen, and particularly to sites more dorsal and lateral than the orbitofrontal cortex (OFC) or ventromedial prefrontal cortex (vmPFC) projections described below (Haber et al., 2006). This set of cortical regions (dorsal anterior cingulate, anterior insula, anterior striatum) is considered the core of the adjacent “Cingulo-opercular” and “Salience” networks, which are critical for a wide variety of cognitive control operations
5. We found functional connections between nucleus accumbens and pregenual medial prefrontal cortex (mPFC), and a separate set with OFC / subgenual cingulate cortex (Fig 4, blue / red). These connections match circuitry known to be critical for reward processing / addiction (Haber and Knutson, 2010; Koob and Volkow, 2016) and mood disorders (Drevets et al., 2008), respectively. The nucleus accumbens is anatomically segregated into a more dorsal shell and a more ventral core (Salgado and Kaplitt, 2015). Here, the nucleus accumbens nodes of these two subnetworks exhibited a similar dorsal/ventral distribution; as such, we tentatively label these nodes the accumbens shell and core, respectively.

Interestingly, these connections only partially converged with connectivity identified in non-human primates, in which the nucleus accumbens core preferentially receives projections from pregenual mPFC, while the shell receives projections from OFC—the reverse of the arrangement we observed here. It is possible that this divergence from non-human primate work is driven by fMRI data quality issues, such as the known presence of susceptibility artifact within ventral prefrontal and orbitofrontal cortex in human fMRI. However, human diffusion tractography imaging has also demonstrated that the accumbens core is more strongly connected to OFC than the shell (Baliki et al., 2013; Cartmell et al., 2019; Xia et al., 2017), convergent with our present findings. We consider an alternate possibility that this divergence represents a true organizational difference between humans and non-human primates related to reward processing.

Several of the connections identified here, including to the posterior and middle putamen, dorsal caudate, and nucleus accumbens core, were restricted to a single target subcortical structure. This represents an increase in target specificity relative to the (Haber, 2016) model, in which similar fronto-striatal projections span the internal capsule (Fig 4A,B, green and orange) or extend out of nucleus accumbens into caudate (Fig 4A,B, red). We consider it most likely that frontal projections to multiple striatal structures do exist in humans, but that projections to secondary structures do not drive striatal function as strongly as projections to the primary target structure.

### Two Cortico-Striatal Subnetworks Diverged from Primate Models but Converged with Human Language Networks

We observed two subnetworks that did not converge with proposed models of non-human primate fronto-striatal connectivity (Haber, 2016). These consisted of two adjacent sets of distributed fronto-temporal regions connected to anterior medial putamen and anterior lateral caudate (Fig 4B, pink / dark green). Based on several lines of evidence, we argue that these connections represent circuitry for language processing. First, the cortical elements of these connections are highly convergent with previous descriptions of Language networks in group-average (Friederici and Gierhan, 2013; Ji et al., 2019) and individual (Braga et al., 2020) human neuroimaging data. Second, as with human language function (Corballis, 2012; Friederici, 2011; Ocklenburg et al., 2014), connections of these two subnetworks were consistently lateralized, with greater extent in the left hemisphere than in the right hemisphere in both cortex and striatum. No other cortico-striatal network structure exhibited consistent lateralization across subjects. Third, the striatal elements of these connections converge well with previous work reporting activation in anterior medial putamen (Friederici et al., 2003) and anterior lateral caudate (Mestres-Missé et al., 2012) in response to specifically effortful, non-automatic language processing. These findings suggest that the striatal elements of these language subnetworks may be critical for learning and executing new language-related tasks, similar to the learning and optimization role ascribed to the striatum in other behavioral domains (Cohen and Frank, 2009).

Some connections of this human cortico-striatal language network do exist in non-human primates. Lateral prefrontal cortex and superior temporal sulcus have been shown to project to each other (Yeterian et al., 2012), and superior temporal cortex projects to lateral caudate and medial putamen (Selemon and Goldman-Rakic, 1985; Yeterian and Pandya, 1998). However, the multiple elements of the language network (inferior frontal gyrus, posterior middle frontal gyrus, medial superior frontal gyrus, superior temporal sulcus, and striatum) are not known to be all interconnected in non-human primates, as we demonstrate here in humans. This supports the idea that the human language network is an evolutionarily novel network scaffolded in part upon existing circuitry present in pre-human primates (Dick and Tremblay, 2012; Friederici, 2017).

### Fronto-striatal Connections Are Organized as a Stepped, not Continuous, Gradient

Prior work has demonstrated that fronto-striatal connections are organized as mirrored cortical and striatal gradients spanning from primary motor cortex / posterior putamen to nucleus accumbens / ventromedial prefrontal cortex. This finding is consistent across humans and non-human primates, and across methodologies ranging from invasive tract-tracing to diffusion MRI to RSFC to task-based fMRI (Haber, 2003; Jarbo and Verstynen, 2015; Marquand et al., 2017; Mestres-Missé et al., 2012; O’Rawe et al., 2019; Raut et al., 2020; Steiner and Tseng, 2010; Vogelsang and D’Esposito, 2018), and similar mirrored gradients have even been observed in mice (Peters et al., 2021). Interestingly, the rostral-caudal orientation of these fronto-striatal gradients closely match—and likely exist within—a proposed whole-brain organizational framework in which multiple mirrored gradients span from unimodal to transmodal regions (Huntenburg et al., 2018; Margulies et al., 2016; Raut et al., 2020; Zhang et al., 2019).

Fronto-striatal gradients are often presented, for visual or mathematical convenience, as if they were smooth and continuous. However, the concept of a smooth, continuous fronto-striatal gradient conflicts with evidence that the brain is organized as discrete, discontinuous areas networked together (Amunts and Zilles, 2015; Brodmann, 1909; Churchland and Sejnowski, 1988; Felleman and Van Essen, 1991; Glasser et al., 2016a; Sejnowski and Churchland, 1989). These areas are established early in the brain’s development, when multiple continuous, overlapping gradients of morphogens and signaling molecules, combined with differential thalamic inputs, induce a process of discrete arealization (O’Leary et al., 2007). While continuous topographic gradients can be present within a given area (e.g. the visuotopic organization of primary visual cortex), these small-scale gradients are discontinuous (and sometimes even reversed) at areal boundaries (Baumann et al., 2013; Felleman and Van Essen, 1991; Leaver and Rauschecker, 2016; Sereno et al., 1995).

Importantly, a progressive rostral-to-caudal series of such discontinuous cortical areas with topographically ordered striatal projections—a stepped gradient—can visually (Fig 5A) and statistically (Fig 6A,B) appear continuous, particularly if the categorical divisions are unknown. Here, direct comparison of continuous gradient models against discontinuous, stepped gradient models demonstrated that the stepped gradient models explained the data much better (Fig 6). This indicates that there is little evidence for a smooth, continuous cross-areal gradient, either as a primary model of human fronto-striatal organization or as a secondary effect superimposed on top of a stepped gradient. Instead, the data is best explained as a stepped rostral-caudal gradient composed of a progressive series of discrete fronto-striatal subnetworks. Interestingly, the explanatory power of the stepped gradient model was particularly strong for connections originating from medial frontal cortex, where it explained almost 2/3rds of the variance in the connection’s location. This suggests that medial frontal projections are more strictly ordered than lateral frontal projections—an intriguing functional dissociation that should be examined in future work.

These results do not invalidate previous findings of a rostral-caudal organization of human fronto-striatal circuitry; rather, they modify the interpretation and implications of those findings. For example, findings that the rostral-caudal organization is related to separable types of goal-directed behaviors (Marquand et al., 2017), and that it represents a hierarchy of information processing, such that more rostral fronto-striatal regions process progressively more complex or abstract information (Choi et al., 2018; Jeon et al., 2014; Nee and Brown, 2013), are consistent with the present observations. However, these prior findings should be interpreted in the context of discrete networked structures that enable those different behaviors and levels of processing, rather than as a continuous gradient. It is possible that the large-scale unimodal-transmodal gradients previously identified across the cortex and subcortex (Huntenburg et al., 2018; Margulies et al., 2016; O’Rawe et al., 2019; Raut et al., 2020; Zhang et al., 2019) may also be best conceptualized as stepped rather than continuous gradients, composed of a series of mirrored network and subnetwork connections.

The present approach for comparing continuous and stepped gradient models does have limitations. First, the infomap technique, while excellent at recovering true networked connections, is not always stable in determining when subnetworks should be combined or split. As a result, the categorical factor entered into the regression model may sometimes have too many or too few categories. Second, using the rostral-caudal location of the strongest functional connection as a dependent variable ignores the possibility that even very strong cortico-striatal connections are many-to-one (as is suggested by the identified subnetworks). Thus, the model may not accurately represent the brain’s true connectional organization. Importantly, both of these issues would be expected to reduce the variance explained of the stepped gradient model, but would not affect the continuous gradient model. The fact that the stepped model outperforms the gradient model despite these limitations is strong evidence that it more accurately represents the organization of fronto-striatal connections.

### Towards a Complete Description of Cortico-Striato-Thalamic Circuits and their Cross-Circuit Integration

It is well established in rodents and non-human primates that cortico-striatal circuits are further projected to the thalamus, and then back to cortex to form feedback loops (Haber, 2003; Nakano et al., 2000). Thus, in principle the cortico-striatal subnetwork structures we observe here should also have representation in the thalamus. While we did observe this representation to some degree, the thalamus did not consistently contain representations of all cortico-striatal subnetworks; and further, the thalamus tended to have extensive representation of subnetworks characterized as noise-related. This suggests that the quality of the MSC fMRI data, while adequate for winner-take-all style whole-network mapping in thalamus (Greene et al., 2020), may not be adequate for the data-driven mapping of highly specific cortico-striato-thalamo-cortical circuits in thalamus. Future work may be able to more broadly conduct such mapping using such higher-quality data collected with advanced multiband sequences.

While the presented delineations of cortico-striatal circuits are neurobiologically compelling and individual-specific, they are incomplete. In examining only the very strongest functional connections to each striatal voxel, we focus on the likely driving inputs to the striatum, but we also exclude the wealth of multi-lobe, modulatory inputs to striatum that have been described in humans (Choi et al., 2012; Greene et al., 2020, 2014; Parkes et al., 2017) and non-human primates (Selemon and Goldman-Rakic, 1985; Yeterian and Van Hoesen, 1978). The current results also represent these circuits as if they are fully segregated from each other. However, non-human primate studies have demonstrated that fronto-striatal circuits are only partially segregated, with some degree of cross-circuit connections (Haber, 2003). Similarly, at the large-scale network level, human RSFC work has also demonstrated cross-network integration zones (Greene et al., 2020). In reproducing analyses from (Greene et al., 2020) at the subnetwork level rather than the network level, we similarly observed extensive cross-subnetwork connections (Fig S4). Untangling the substantial complexity of these multi-subnetwork connections for comprehensive evaluation and categorization must be conducted in future work.

### Mapping Discrete Cortico-Striatal Circuits in Individual Humans

In describing cortico-striatal organization as a stepped gradient composed of discrete subnetworks, the present results have important implications for the future study of the human striatum. The discrete subnetworks present within the striatum likely enable a variety of different cognitive and motor processes that can be localized to each separate, individual-specific striatal network structure. With such localization, we can generate strong hypotheses about how individual-specific symptoms observed in neurosychiatric disorders might arise from disruption of those specific fronto-striatal circuits, or about how neurodegenerative diseases or brain injuries that damage specific fronto-striatal circuits would impair associated cognitive and motor functions. We can then explore whether symptoms can be alleviated or impaired function restored via brain stimulation approaches, including noninvasive TMS targeting the cortical sources of these fronto-striatal projections, or deep electrode implantation targeting their striatal targets.

## METHODS

### Subjects

Data were collected from ten healthy, right-handed, young adult subjects (5 females; age: 24-34). Two of the subjects are authors (NUFD and SMN), and the remaining subjects were recruited from the Washington University community. Informed consent was obtained from all participants. The study was approved by the Washington University School of Medicine Human Studies Committee and Institutional Review Board. Other findings using these participants have been previously reported in (Gordon et al., 2017c; Gratton et al., 2018; Greene et al., 2020; Marek et al., 2018; Sylvester et al., 2020).

### MRI image acquisition

Imaging for each subject was performed on a Siemens TRIO 3T MRI scanner over the course of 12 sessions conducted on separate days, each beginning at midnight. Structural MRI was conducted across two separate days. In total, four T1-weighted images (sagittal, 224 slices, 0.8 mm isotropic resolution, TE=3.74 ms, TR=2400 ms, TI=1000 ms, flip angle = 8 degrees), four T2-weighted images (sagittal, 224 slices, 0.8 mm isotropic resolution, TE=479 ms, TR=3200 ms), four MRA (transverse, 0.6 x 0.6, x 1.0mm, 44 slices, TR=25ms, TE=3.34ms) and eight MRVs, including four in coronal and four in sagittal orientations (sagittal: 0.8 x 0.8 x 2.0mm thickness, 120 slices, TR=27ms, TE=7.05ms; coronal: 0.7 x 0.7 x 2.5mm thickness, 128 slices, TR=28ms TE= 7.18ms), were obtained for each subject. Analyses of the MRA and MRV scans are not reported here.

On ten subsequent days, each subject underwent 1.5 hours of functional MRI scanning beginning at midnight. In each session, we first collected thirty contiguous minutes of resting state fMRI data, in which subjects visually fixated on a white crosshair presented against a black background. Each subject was then scanned during performance of three separate tasks, which are not examined here. Across all sessions, each subject was scanned for 300 total minutes during the resting state. All functional imaging was performed using a gradient-echo EPI sequence (TR = 2.2 s, TE = 27 ms, flip angle = 90°, voxel size = 4 mm x 4 mm x 4 mm, 36 slices). In each session, one gradient echo field map sequence was acquired with the same prescription as the functional images. An EyeLink 1000 eye-tracking system (http://www.sr-research.com) allowed continuous monitoring of subjects’ eyes in order to check for periods of prolonged eye closure, potentially indicating sleep. Only one subject (MSC08) demonstrated prolonged eye closures.

### Cortical surface generation

Generation of cortical surfaces from the MRI data followed a procedure similar to that previously described in (Marek et al., 2018). First, anatomical surfaces were generated from the subject’s average T1-weighted image in native volumetric space using FreeSurfer’s default recon-all processing pipeline (version 5.3). This pipeline first conducted brain extraction and segmentation. After this step, segmentations were hand-edited to maximize accuracy. Subsequently, the remainder of the recon-all pipeline was conducted on the hand-edited segmentations, including generation of white matter and pial surfaces, inflation of the surfaces to a sphere, and surface shape-based spherical registration of the subject’s original surface to the fsaverage surface (Dale et al., 1999; Fischl et al., 1999). The fsaverage-registered left and right hemisphere surfaces were brought into register with each other using deformation maps from a landmark-based registration of left and right fsaverage surfaces to a hybrid left-right fsaverage surface (‘fs_LR’; Van Essen et al., 2012). These fs_LR spherical template meshes were input to a flexible Multi-modal surface Matching (MSM) algorithm using sulc features to register templates to the atlas mesh (Robinson et al., 2014). These newly registered surfaces were then down-sampled to a 32,492 vertex surface (fs_LR 32k) for each hemisphere. The various structural metric data (thickness, curvature, etc.) from the original surfaces to the fs_LR 32k surface were composed into a single deformation map allowing for one step resampling. MSM registration provided a more optimal fit of pial and white surfaces and reduced areal distortion (Glasser et al., 2016b). These various surfaces in native stereotaxic space were then transformed into atlas space (711-2B) by applying the previously calculated T1-to-atlas transformation.

### fMRI Preprocessing

Functional data were preprocessed to reduce artifacts and to maximize cross-session registration. All sessions underwent correction of odd vs. even slice intensity differences attributable to interleaved acquisition, intensity normalization to a whole brain mode value of 1000, and within run correction for head movement. Atlas transformation was computed by registering the mean intensity image from a single BOLD session to Talairach atlas space (Talairach and Tournoux, 1988) via the average high-resolution T2-weighted image and average high-resolution T1-weighted image. All subsequent BOLD sessions were linearly registered to this first session. This atlas transformation, mean field distortion correction (see below), and resampling to 2-mm isotropic atlas space were combined into a single interpolation using FSL’s applywarp tool (Smith et al., 2004). All subsequent operations were performed on the atlas-transformed volumetric time series.

### Distortion correction

A mean field map was generated based on the field maps collected in each subject (Laumann et al., 2015). This mean field map was then linearly registered to each session and applied to that session for distortion correction. To generate the mean field map the following procedure was used: (1) Field map magnitude images were mutually co-registered. (2) Transforms between all sessions were resolved. Transform resolution reconstructs the n-1 transforms between all images using the n(n-1)/2 computed transform pairs. (3) The resolved transforms were applied to generate a mean magnitude image. (4) The mean magnitude image was registered to an atlas representative template. (5) Individual session magnitude image to atlas space transforms were computed by composing the session-to-mean and mean-to-atlas transforms. (6) Phase images were then transformed to atlas space using the composed transforms, and a mean phase image in atlas space was computed.

Application of mean field map to individual fMRI sessions: (1) For each session, field map uncorrected data were registered to atlas space, as above. (2) The generated transformation matrix was then inverted and applied to the mean field map to bring the mean field map into the session space. (3) The mean field map was used to correct distortion in each native-space run of resting state and task data in the session. (4) The undistorted data were then re-registered to atlas space. (5) This new transformation matrix and the mean field map then were applied together to resample each run of resting state and task data in the session to undistorted atlas space in a single step.

### RSFC Preprocessing

Additional preprocessing steps to reduce spurious variance unlikely to reflect neuronal activity were executed as recommended in (Ciric et al., 2017; Power et al., 2014). First, temporal masks were created to flag motion-contaminated frames. We observed that two subjects (MSC 03 and MSC 10) had a high-frequency artifact in the motion estimates calculated in the phase encode (anterior-posterior) direction that did not appear to reflect biological movement. We thus filtered the motion estimate time courses in this direction only to retain effects occurring below 0.1 Hz in all subjects for consistency (Gratton et al., 2019). Motion contaminated volumes were then identified by frame-by-frame displacement (FD). Frames with FD > 0.2mm were flagged as motion-contaminated. Across all subjects, these masks censored 28% ± 18% (range: 6% – 67%) of the data; on average, subjects retained 5929 ± 1508 volumes (range: 2733 – 7667), corresponding to 217 ± 55 minutes (range: 100 – 281).

After computing the temporal masks for high motion frame censoring, the data were processed with the following steps: (i) demeaning and detrending, (ii) linear interpolation across censored frames using so that continuous data can be passed through (iii) a band-pass filter (0.005 Hz < f < 0.01 Hz) without re-introducing nuisance signals (Hallquist et al., 2013) or contaminating frames near high motion frames (Carp, 2013).

Next, the filtered BOLD time series underwent a component-based nuisance regression approach (Marek et al., 2018). Nuisance regression using time series extracted from white matter and cerebrospinal fluid (CSF) assumes that variance in such regions is unlikely to reflect neural activity. Variance in these regions is known to correspond largely to physiological noise (e.g., CSF pulsations), arterial pCO2-dependent changes in T2*-weighted intensity and motion artifact; this spurious variance is widely shared with regions of interest in gray matter. We also included the mean signal averaged over the whole brain as a nuisance regressor. Global signal regression (GSR) has been controversial. However, the available evidence indicates that GSR is a highly effective de-noising strategy (Ciric et al., 2017; Power et al., 2015).

Nuisance regressors were extracted from white matter and ventricle masks, first segmented by FreeSurfer (Fischl, 2012), then spatially resampled in register with the fMRI data. Voxels surrounding the edge of the brain are particularly susceptible to motion artifacts and CSF pulsations (Patriat et al., 2015; Satterthwaite et al., 2013); hence, a third nuisance mask was created for the extra-axial compartment by thresholding the temporal standard deviation image (SDt > 2.5%), excluding a dilated whole brain mask. Voxel-wise nuisance time series were dimensionality reduced as in CompCor (Behzadi et al., 2007), except that the number of retained regressors, rather than being a fixed quantity, was determined, for each noise compartment, by orthogonalization of the covariance matrix and retaining components ordered by decreasing eigenvalue up to a condition number of 30 (max eigenvalue / min eigenvalue > 30). The retained components across all compartments formed the columns of a design matrix, X, along with the global signal, its first derivative, and the six time series derived by retrospective motion correction. The columns of X are likely to exhibit substantial co-linearity. Therefore, to prevent numerical instability owing to rank-deficiency during nuisance regression, a second-level SVD was applied to XX^T^ to impose an upper limit of 250 on the condition number. This final set of regressors was applied in a single step to the filtered, interpolated BOLD time series, with censored data ignored during beta estimation. Finally, the data were upsampled to 2mm isotropic voxels. Censored frames were then excised from the data for all subsequent analyses.

### Surface processing and CIFTI generation of BOLD data

Surface processing of BOLD data proceeded through the following steps. First, the BOLD fMRI volumetric timeseries were sampled to each subject’s original mid-thickness left and right-hemisphere surfaces (generated as the average of the white and pial surfaces) using the ribbon-constrained sampling procedure available in Connectome Workbench 1.0. This procedure samples data from voxels within the gray matter ribbon (i.e., between the white and pial surfaces) that lie in a cylinder orthogonal to the local mid-thickness surface weighted by the extent to which the voxel falls within the ribbon. Voxels with a timeseries coefficient of variation 0.5 standard deviations higher than the mean coefficient of variation of nearby voxels (within a 5 mm sigma Gaussian neighborhood) were excluded from the volume to surface sampling, as described in (Glasser et al., 2013). Once sampled to the surface, timecourses were deformed and resampled from the individual’s original surface to the 32k fs_LR surface in a single step using the deformation map generated above (in “Cortical surface generation”). This resampling allows point-to-point comparison between each individual registered to this surface space.

These surfaces were then combined with volumetric subcortical and cerebellar data into the CIFTI format using Connectome Workbench (Marcus et al., 2011), creating full brain timecourses excluding non-gray matter tissue. Subcortical (including accumbens, amygdala, caudate, hippocampus, pallidum, putamen, and thalamus) and cerebellar voxels were selected based on the FreeSurfer segmentation of the individual subject’s native-space average T1, transformed into atlas space, and manually inspected. Finally, the BOLD timecourses were smoothed with a geodesic 2D (for surface data) or Euclidean 3D (for volumetric data) Gaussian kernel of σ = 2.55 mm.

### Regression of adjacent cortical tissue from RSFC BOLD

Some striatal regions, particularly including lateral putamen, are in close anatomical proximity to cortex, resulting in spurious functional coupling between the cortical vertices and adjacent subcortical voxels. To reduce this artifact, RSFC BOLD time series from all vertices falling within 20mm Euclidean distance of a source voxel were averaged and then regressed from the voxel time series (Buckner et al., 2011; Greene et al., 2020; Marek et al., 2018). The resulting residual timeseries were used for all subsequent analyses.

### Mapping multi-scale network structure

The network organization of each subject’s brain was delineated following (Gordon et al., 2020) using the graph-theory-based Infomap algorithm for community detection (Rosvall and Bergstrom, 2008). In this approach, we calculated the cross-correlation matrix of the time courses from all brain vertices (on the cortical surfaces) and voxels (in subcortical structures), concatenated across sessions. Correlations between vertices/voxels within 30 mm of each other were set to zero in this matrix to avoid basing network membership on correlations attributable to spatial smoothing. Geodesic distance was used for within-hemisphere surface connections and Euclidean distance for subcortical-to-cortical connections. Connections between subcortical structures were disallowed, as we observed extremely high connectivities within nearly the entire basal ganglia that would prevent network structures from emerging. Inter-hemispheric connections between the cortical surfaces were retained, as smoothing was not performed across the mid-sagittal plane.

We observed that connectivity patterns within regions known to have low BOLD signal due to susceptibility artifact dropout (e.g. ventral anterior temporal lobe, portions of orbitofrontal cortex) were effectively random. To avoid having the delineated network structures distorted by such random connections, we first calculated a set of common low-signal regions as the vertices in which the average mode-1000 normalized BOLD signal across subjects and timepoints was less than 750 (as in Gordon et al., 2016; Wig et al., 2014). All connections to these low-signal regions were set to zero.

This matrix was then thresholded to retain at least the strongest .1% of connections to each vertex and voxel, following observations in (Gordon et al., 2020) that this level of network division represents divisions that, across these subjects, optimally divide the resting state data and explain task activation patterns.

These thresholded matrices were used as inputs for the Infomap algorithm, which calculated community assignments (representing brain network structures) separately for each threshold. Small networks with 10 or fewer vertices / voxels were considered unassigned and removed from further consideration. The above analysis was conducted in each individual subject.

### Identifying matched striatal subnetworks in individuals

For each subject, we considered only subnetworks that had at least some representation in the striatum (the individually defined caudate, putamen, nucleus accumbens, and globus pallidus). Following (Gordon et al., 2020), we visually examined the cortical and subcortical topographies of each subnetwork with striatal representation, as well as their topological arrangement relative to each other. After careful consideration of the subnetworks observable in this population, matched subnetworks were identified based on the following heuristic rules:

A subnetwork in inferior ventral striatum usually also had representation in subgenual cingulate and/or orbitofrontal cortex.

A subnetwork in anterior ventral striatum had representation in bilateral pregenual cingulate cortex. This subnetwork was previously described in (Gordon et al., 2020).

A subnetwork in superior ventral striatum had representation in anterior cingulate cortex and middle insula.

A subnetwork that was spatially variable within striatum, with representation most often in medial putamen, but was consistently present in dorsal anterior cingulate cortex and dorsal insula.

A subnetwork in lateral middle putamen and posterior dorsomedial prefrontal cortex.

A subnetwork in dorsal caudate that had consistent representation as the dorsomedial cluster of the Fronto-Parietal network and inconsistent representation in lateral frontal and parietal cortex.

A subnetwork in medial anterior putamen (especially left putamen) that had consistent cortical representation converging with the Language network (Braga et al., 2020).

A subnetwork in lateral anterior caudate (especially left caudate) that had representation to a set of cortical regions directly anterior to the Language network above.

A subnetwork (sometimes multiple subnetworks) in ventral posterior putamen that had representation somewhere in the central sulcus, with the location of the representation varying substantially by subject.

Additionally, we commonly observed subnetworks that were generally in posterior lateral putamen (but with variable representation across subjects) that had cortical representation only in middle insular cortex, physically adjacent to the striatal representation, or (in one subject) in unusual locations such as medial occipital lobe. These subnetworks also often cut implausibly across disjoint anatomical structures such as putamen and thalamus. These subnetworks were tentatively classified as noise-related and were ignored for future analyses.

### Computing lobe-wise striatal subnetwork representation

For each individual subject, we defined the Frontal, Insular, Parietal, Temporal, and Occipital lobes based on the lobe identities of the regions of the subject-specific Freesurfer-generated Destrieux atlas parcellation, which was deformed into fs_LR_32k space to match the functional data. Lobe-wise representation was computed within each subject as the percent of cortical vertices among all non-noise subnetworks that were within each lobe.

### Visualizing subnetwork overlap across subjects

For each matched non-noise subnetwork, the overlap across individuals was visualized using Connectome Workbench. For cortex, the number of individuals with the subnetwork present was calculated in each vertex. For striatum, this was less straightforward, as the subcortical structures were individualized, and so were different shapes and sizes across people. To represent overlap in striatum, we first mapped the subcortical voxels of each individual among MSC02-10 to the subcortical structures in MSC01. This was done by assigning each voxel in the MSC01 structures the network ID of the nearest (by Euclidean distance) subcortical voxel in that individual. Overlap was then calculated across MSC01-space subject subnetworks as the number of individuals with the subnetwork present in each MSC01-space voxel.

### Defining position along a rostral-caudal axis in striatum and frontal cortex

For each subject, we calculated four separate curves to fit the rostral-caudal orientation of 1) bilateral caudate, 2) left putamen, 3) right putamen, and 4) frontal cortex.

For caudate, the structure is oriented vertically, and the rostral-caudal organization is best represented as a curve running from ventral striatum up through caudate head and back to caudate tail—that is, a curve in the y and z dimensions, with the x dimension irrelevant. We thus created a rostral-caudal axis as a lowess curve fit to the y- and z-coordinates of the caudate voxels. This curve represents the approximate geometric center of the structure in the y and z dimensions. In the dorsal posterior portion of the caudate (body and tail), the rostral-caudal organization is simply anterior-to-posterior. Therefore, once the curve reached the maximum z coordinate of the structure, the lowess fit was terminated and the axis was continued by projecting straight backwards along the y axis. For this fitting, nucleus accumbens was combined with caudate. Additionally, left and right caudate were combined, as the difference between the left and right structures (in the x-axis) was irrelevant for the fit.

For putamen, the structure is oriented horizontally, and thus the rostral-caudal organization is best represented as a curve in the x and y dimensions, with the z dimension irrelevant. Thus, as with caudate, we created a rostral-caudal axis as a lowess curve fitted to the x- and y-coordinates of the putamen voxels, representing the approximate geometric center of the structure in the x and y dimensions. Once the curve reached the maximum x coordinate of the structure, the lowess fit was terminated and the axis was continued by projecting straight backwards along the y axis. Because the left and right putamen are mirrored in the x dimension (rather than oriented equivalently, as with the caudate z dimension), the fit was necessarily conducted separately for the left and right structures. For this analysis, globus pallidus was not included in the fit.

For frontal cortex, the structure is oriented vertically, as with caudate. Therefore, as with caudate, we created a rostral-caudal axis as a lowess curve fit to the y- and z-coordinates of the cortical vertices in the frontal cortex. For each subject, frontal cortex was defined using the subject-specific Destrieux atlas parcellation, as above, except that we also included anterior insula regions, which commonly included substantial representation of striatal networks (excluding insula regions did not change the results; see Fig S5B). As with caudate, we terminated the curve and projected backwards along the y axis once the curve reached the maximum z coordinate.

Finally, we calculated the coordinate of each striatal voxel and each cortical vertex along the rostral-caudal axis. First, the distances along the four rostral-caudal fit curve axes were converted to rank value (rank within each curve) to ensure comparability across structures. Then, for each voxel / vertex, the rostral-caudal coordinate was taken to be the rank of the point on the relevant structure’s rostral-caudal axis closest (i.e. minimum Euclidean distance) to that voxel/vertex. Note that this calculation—the closest point on the relevant axis—is identical to how x/y/z axis coordinates are defined in a standard 3D space.

In each case, rostral-caudal positions were converted to rank order along the rostral-caudal gradient to account for nonuniform effects of position.

### Examining continuous and discrete organizations of strong fronto-striatal connections

We conducted two separate sets of analyses: One in which we identified the most strongly connected striatal target voxel for each frontal cortex source vertex, and the second in which we identified the most strongly connected frontal source vertex for each striatal target voxel.

#### Regression Set 1: Cortex→Striatum

For each frontal cortex vertex, we identified the striatal voxel with which it had the strongest functional connection. This was done by calculating correlations between the timecourses of all subcortical voxels and all frontal cortex vertices (frontal cortex defined as above). For each frontal vertex, the striatal voxel with the most strongly correlated timecourse was taken to be that vertex’s “target”—the voxel most likely to be directly anatomically connected to that voxel. Voxels with correlations to their sources that were weaker than R=.2 were considered too weak to be reliable and were excluded from subsequent analyses. However, inclusion of these voxels does not change the results; see Fig S5A.

We tested which factors of a frontal source vertex explain the rostral-caudal position of its striatal target. We first tested whether fronto-striatal connections could appear to be a continuous rostral-caudal gradient, in which continuous movement from vertex to vertex along the frontal cortex rostral-caudal axis is mirrored by a continuous rostral-caudal movement of those vertices’ striatal targets. Specifically, separately in each subject, we tested whether the rostral-caudal position of cortical vertices was correlated with the rostral-caudal position of their striatal targets. We evaluated this effect as the adjusted square of the correlation value (R^2^), representing the variance in striatal voxel position explained by cortical target position, corrected for the use of the single regressor.

We then tested whether fronto-striatal connections could appear as a stepped gradient composed of discrete networks, in which the location of a striatal target voxel is explained by the network of its frontal source vertex. Specifically, separately in each subject, we conducted a one-way ANOVA with the network identity of frontal vertices as an independent variable and the rostral-caudal position of their striatal targets as the dependent variable. We evaluated this effect as the adjusted R^2^ of the main factor of the ANOVA, representing the variance in striatal target position explained by frontal vertex network ID, corrected for the number of levels in the ANOVA. Note that this adjusted R^2^ value is directly comparable to the adjusted R^2^ value from the correlation above.

Finally, we tested whether fronto-striatal connections could appear as a continuous or a stepped rostral-caudal gradient when accounting for the other organization, as well as which organization explained more variance. Separately in each subject, we conducted a one-way ANCOVA with the network identity of frontal vertices as one discrete independent variable, the rostral-caudal position of frontal vertices as a second, continuous variable, and the rostral-caudal position of the striatal targets as the dependent variable. The explanatory power of each factor (continuous gradient vs discrete network) was compared against each other by calculating and computing the adjusted R^2^ of each factor within the ANCOVA.

#### Regression Set 2: Striatum→Cortex

These regressions were conducted using the same logic as above, but using the striatal voxels as independent variables and the frontal vertices as dependent variables. Specifically, for each striatal voxel, we identified the frontal cortex vertex with which it had the strongest functional connection. This was done as above by calculating correlations between the timecourses of all subcortical voxels and all frontal cortex vertices. For each voxel, the cortical vertex with the most strongly correlated timecourse was taken to be its “source”. Voxels with correlations to their sources that were weaker than R=.2 were excluded from subsequent analyses (again, inclusion of these voxels does not change results; Fig S5A).

We tested which factors of a striatal target voxel explain the rostral-caudal position of its frontal source. We first tested whether the rostral-caudal position of striatal voxels was correlated with the rostral-caudal position of their frontal sources. We evaluated this effect as the adjusted square of the correlation value (R^2^), representing the variance in fontal vertex position explained by striatal voxel position, corrected for the use of the single regressor.

We then tested whether the location of a frontal source vertex is explained by the network of its striatal target voxel by conducting a one-way ANOVA with the network identity of striatal voxels as an independent variable and the rostral-caudal position of their frontal sources as the dependent variable. We evaluated this effect as the adjusted R^2^ of the main factor of the ANOVA, representing the variance in frontal source position explained by striatal voxel network ID, corrected for the number of levels in the ANOVA. This adjusted R^2^ value is directly comparable to the adjusted R^2^ value from the correlation above.

Finally, we tested whether fronto-striatal connections could appear as a continuous or a stepped rostral-caudal gradient when accounting for the other organization. Separately in each subject, we conducted a one-way ANCOVA with the network identity of striatal voxels as one discrete independent variable, the rostral-caudal position of striatal voxels as a second, continuous variable, and the rostral-caudal position of the frontal source vertices as the dependent variable. The explanatory power of each factor was compared against each other by calculating and computing the adjusted R^2^ of each factor within the ANCOVA.

## DATA AND SOFTWARE AVAILABILITY

### Data

Raw MRI data from the MSC Dataset, as well as segmented cortical surfaces, preprocessed volumetric and cifti-space RSFC timecourses, and preprocessed task timecourses and contrasts, have been deposited in the Openneuro data repository (https://openneuro.org/datasets/ds000224/versions/1.0.2) under the label “Midnight Scan Club”.

### Code

Code to perform all preprocessing and analysis is available at https://github.com/MidnightScanClub.

## Acknowledgements

This work was supported by NIH grants F31NS110332 (D.J.N.), NS088590 (N.U.F.D.), TR000448 (N.U.F.D.), MH1000872 (T.O.L.), 1R25MH112473 (T.O.L.), 5T32 MH100019-02 (S.M.), MH104592 (D.J.G.), 1P30NS098577 (to the Neuroimaging Informatics and Analysis Center); Kiwanis Neuroscience Research Foundation (N.U.F.D. and B.L.S.); the Jacobs Foundation grant 2016121703 (N.U.F.D.); the Child Neurology Foundation (N.U.F.D.); the McDonnell Center for Systems Neuroscience (N.U.F.D. and B.L.S.); the Mallinckrodt Institute of Radiology grant 14-011 (N.U.F.D.); the Hope Center for Neurological Disorders (N.U.F.D., B.L.S., and S.E.P.). The views expressed in this article are those of the authors and do not necessarily reflect the position or policy of the Department of Veterans Affairs or the U.S. government.

## Contributions

DJG, SMN, JMH, AMN, and NUFD collected the data. EMG, TOL, SMN, DJG, and AZS processed and analyzed the data. EMG, SMN, and NUFD wrote the paper with input from all authors.

---

**Table S1:** For each identified subnetwork, description of locations of common appearance within striatum and cortex, as well as which large-scale network the subnetwork was most commonly contained within.

| Color in Figures | Striatal representation | Cortical representation | Mode large-scale network | Notes |
| --- | --- | --- | --- | --- |
| Magenta | Posterior putamen | Variable locations in central sulcus | Motor |  |
| Light green | Lateral middle putamen | Posterior dorsomedial prefrontal cortex | Cingulo-opercular |  |
| Pink | Anterior medial putamen, sometimes lateral caudate head | Dorsomedial prefrontal, inferior frontal (pars opercularis and/or orbitalis), posterior middle frontal, superior temporal | Language | Lateralized; cortical distribution looks like language network |
| Yellow | Medial caudate head, anterior medial putamen | Dorsomedial prefrontal cortex, anterior middle insula | Fronto-parietal | Cortical representations highly punctate |
| Brown | Anterior globus pallidus and anterior or medial putamen | Dorsal anterior cingulate<br>Dorsal anterior insula | Cingulo-opercular | Striatal representations generally have poor overlap |
| Purple | Ventral caudate and/or anterior ventral putamen | Anterior cingulate<br>Middle insula | Salience |  |
| Light Blue | Anterior medial ventral striatum | Pregenual cingulate<br>Ventral anterior insula (8/10) | Salience |  |
| Dark red | Inferior nucleus accumbens | Orbitofrontal cortex (8/10 subs)<br>Subgenual cingulate (6/10) | Default | Subgenual cluster sometimes very small |
| Dark green | Lateral anterior caudate, medial to location of medial putamen subnetwork above | Adjacent to location of medial putamen subnetwork above | Default | Not found in 2/10 subjects because it was incorporated into medial putamen subnetwork above |
| Orange | Dorsal caudate | Dorsomedial prefrontal cortex<br>Lateral prefrontal cortex | Fronto-parietal | Lateral prefrontal representation punctate in 4/10 subjects, very extensive in 6/10 |
| Dark gray | Inconsistent regions in posterior lateral putamen; often cut implausibly across disjoint structures such as putamen and thalamus | Inconsistent regions in middle insula (adjacent to putamen representation), or, in 1/10 subjects, occipital lobe | N/A | Interpreted as noise-related |

**Figure S1:**
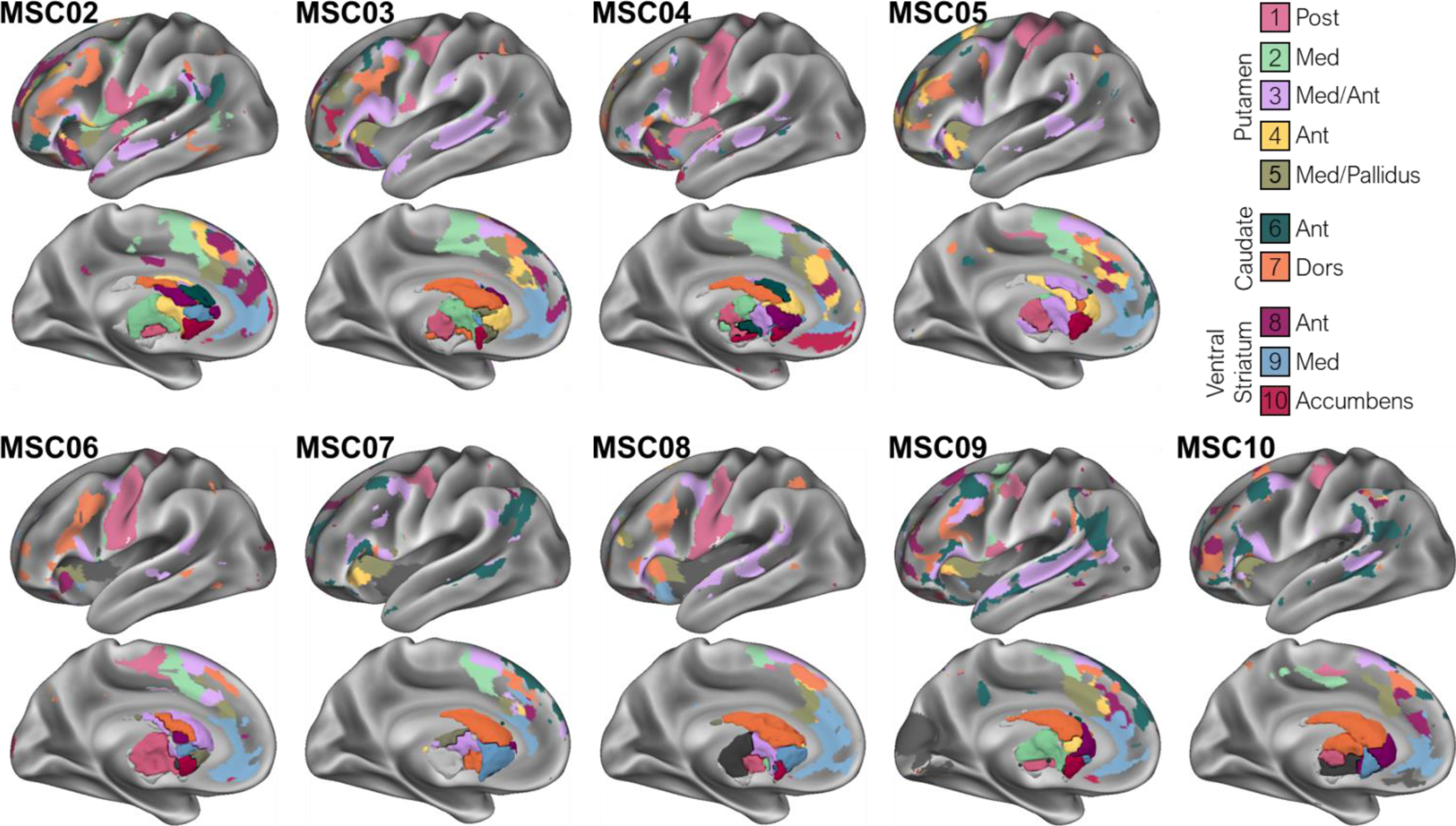
Representation of matched striatal subnetworks in subjects MSC02-10 within left hemisphere cortex and striatum.

**Figure S2:**
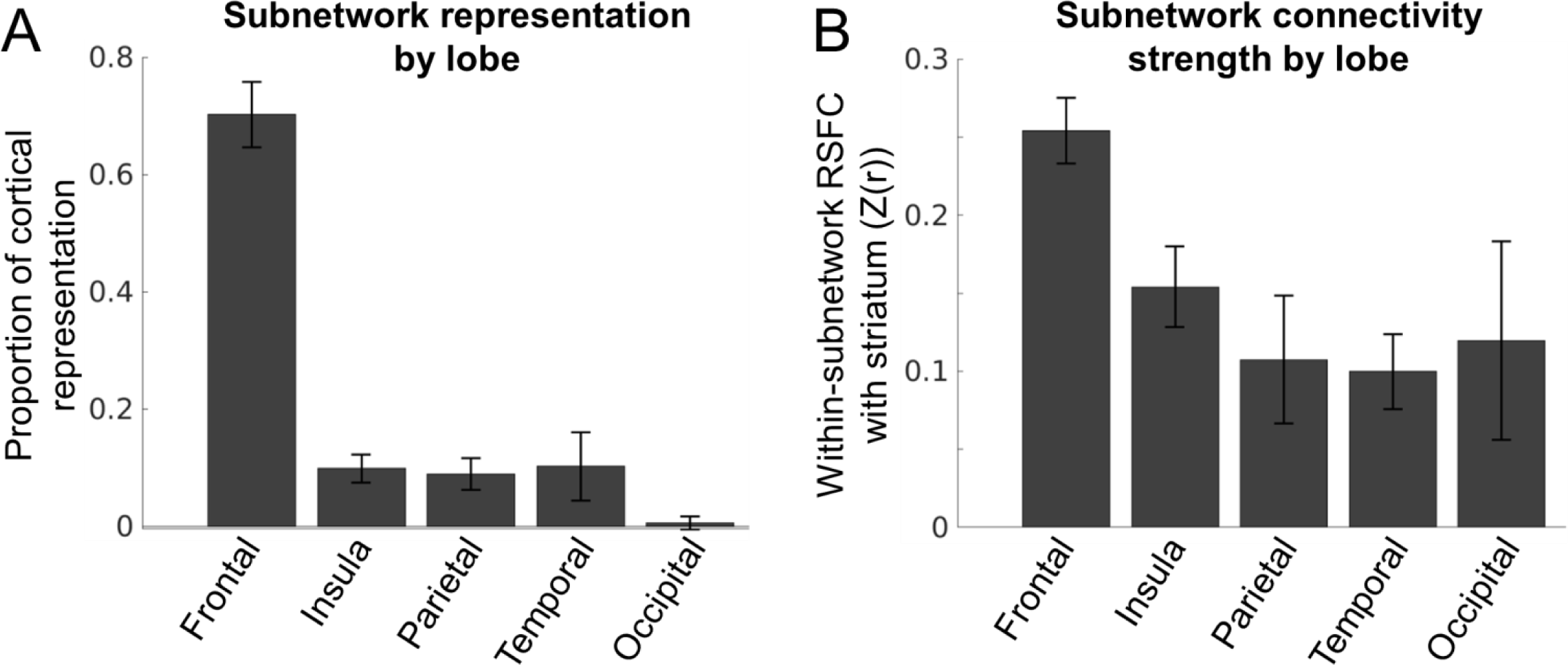
Striatal subnetworks are primarily connected to frontal cortex. A) Percent of cortical subnetwork vertices present within each cortical lobe, averaged across subnetworks and subjects. B) Strength of within-subnetwork functional connectivity between striatum and the vertices within each cortical lobe, averaged across subnetworks and subjects. Error bars represent standard deviation across subjects.

**Figure S3:**
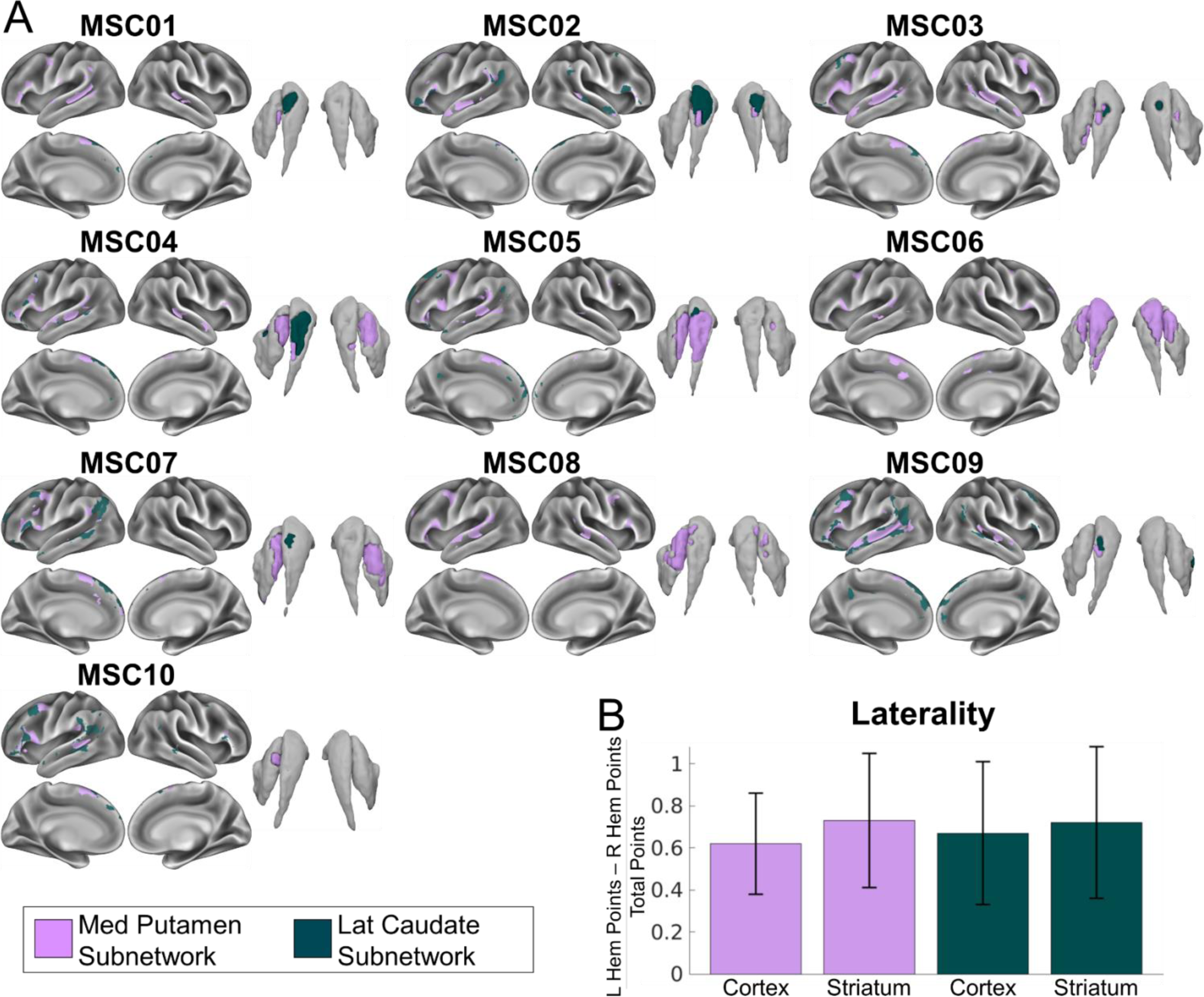
Lateralized cortico-striatal subnetworks consistent with human language networks. A) In each subject, bilateral cortical and striatal representations of two subnetworks consistent with known human language networks. B) Both subnetworks were lateralized (with greater representation in left than right hemisphere) in both cortex and striatum in every subject. No other subnetwork was consistently lateralized across subjects.

**Figure S4:**
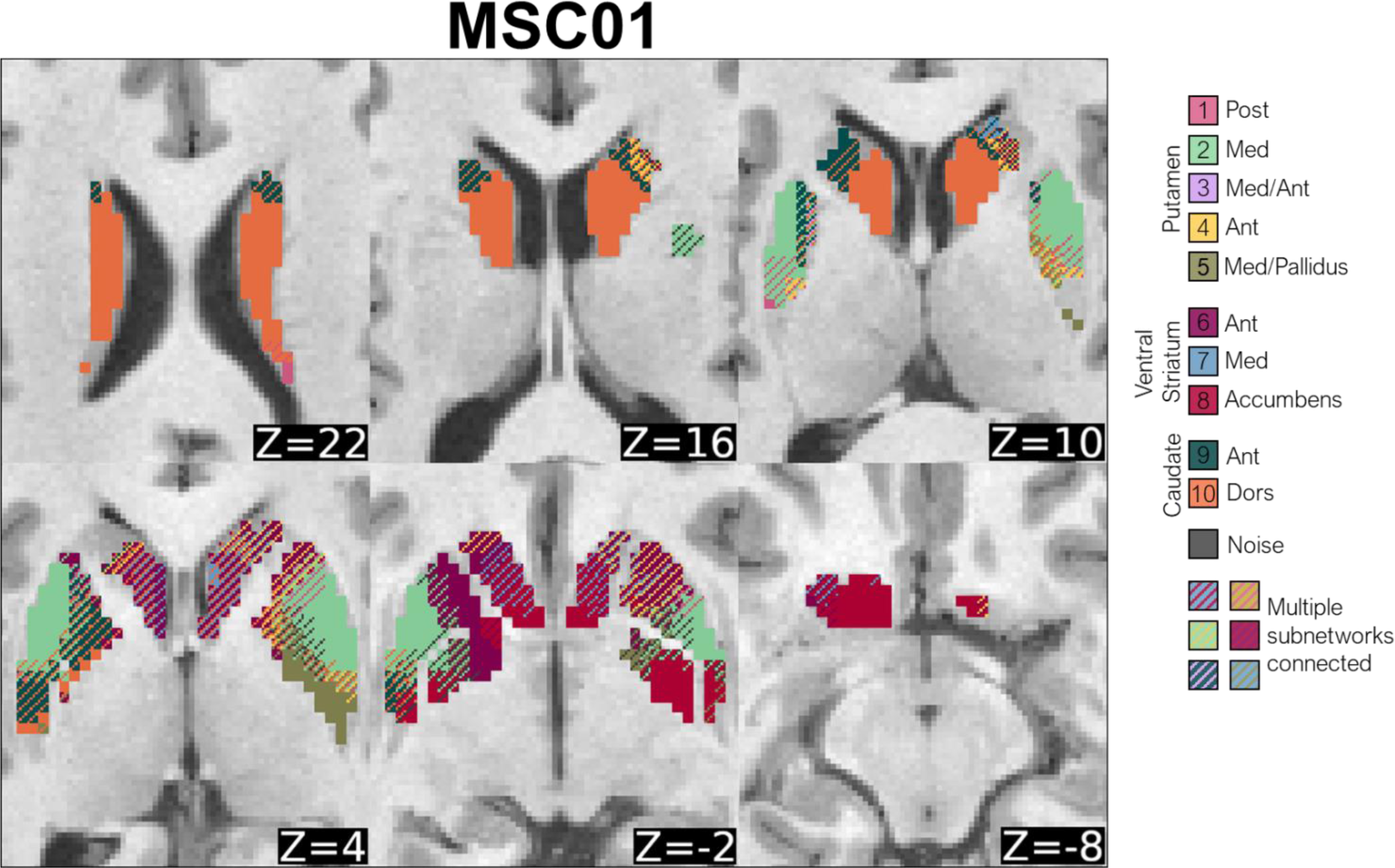
Cross-subnetwork integration in subject MSC01. Voxels with preferential RSFC to one network (network-specific) are represented by solid colors, and voxels functionally connected to multiple networks (integrative) are represented by cross-hatching. Analyses performed following procedures in Greene et al. (2020), except that we evaluated voxel RSFC to matched fronto-striatal subjects rather than to whole networks.

**Figure S5:**
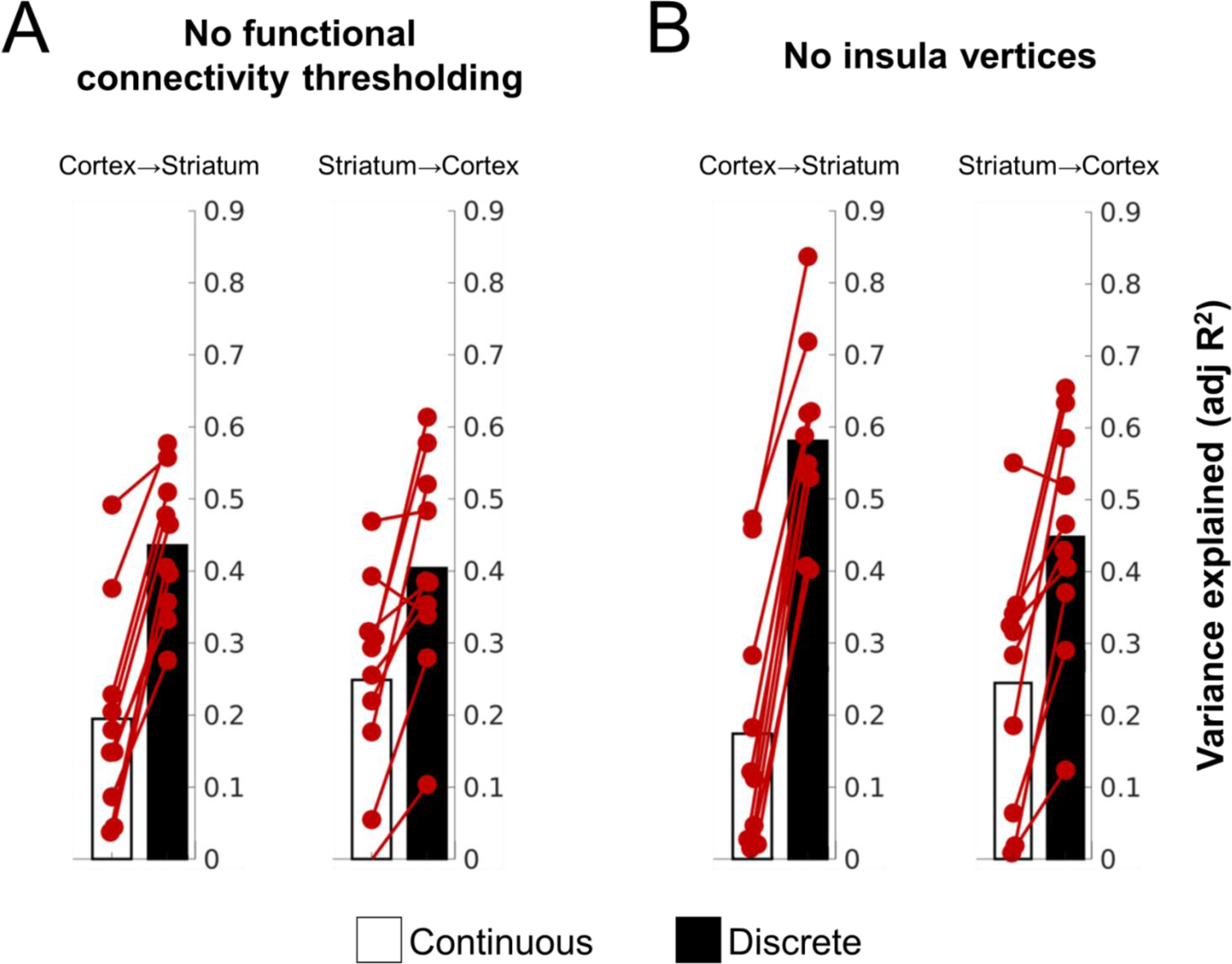
Fronto-striatal organization explained by continuous and stepped gradients after alternate analysis procedures. A) With no minimum RSFC thresholding (minimum RSFC set to Z(r)=.2 in main text analyses), discrete network identity explained more variance than continuous rostral-caudal position for both the Cortex→Striatum (left) and the Striatum→Cortex (right) analyses. B) After excluding anterior insula voxels from consideration, discrete network identity explained more variance than continuous rostral-caudal position for both the Cortex→Striatum (left) and the Striatum→Cortex (right) analyses.

